# Phase-dependent stimulation response is shaped by the brain’s transient collective dynamics

**DOI:** 10.1101/2023.05.09.539965

**Authors:** Sophie Benitez Stulz, Boris Gutkin, Matthieu Gilson, Demian Battaglia

## Abstract

Exogenous stimulation is a promising tool for investigating and altering cognitive processes in the brain, with potential clinical applications. Following experimental observations, we hypothesise that the effect of stimulation crucially depends on the endogenous dynamics of the brain. Our study explores how local and global dynamical properties, like the stimulation phase of regional oscillatory activity and the transient network states, modulate the effect of single pulse stimulation in a large-scale network. Our findings demonstrate that the effect of stimulation strongly depends on the interplay between stimulated phase, transient network state, and brain region. Importantly, we show that stimulation is not only state-dependent but can also induce global state switching. Lastly, predicting the effect of stimulation by using machine learning shows that state-aware measures can increase the performance by up to 40%. Our results suggest that a fine characterisation of the complex brain dynamics in experimental setups is essential for improving the reliability of exogenous stimulation.

---

Brain stimulation has evolved into a promising tool for investigating neurotypical and atypical processes as well as treating pathological processes (*1–6*). While the range of possible applications steadily increases, stimulation outcomes do not always reach the necessary reliability and instead exhibit large intra and inter-subject variability (*7–10*). This variability cannot exclusively be attributed to the used stimulation methodology, but could also reflect the underlying endogenous dynamics of the brain (*11–14*). For instance, several studies showed a phase-dependence in the efficacy of stimulation, with respect to local neuronal activity (*15*– *21*), as well as in responses to perceptual stimuli (*22–24*). This empirical phenomenon is backed by theoretical arguments from complex network systems (*25–27*). Phase, however, is not the only aspect of dynamics that can affect stimulation outcomes. Transient activity states (e.g. EEG microstates) may further modulate the effect of stimulation (*28*). In return, stimulation may modify the dynamical state at various spatio-temporal scales (*29–32*), revealing complex mutual interdependencies which are difficult to disentangle.

To understand how transient endogenous dynamics interacts with exogenous stimulation, we use a large-scale brain model displaying complex oscillatory dynamics, thus overcoming the limitations imposed by the noisiness and the scarcity of empirical data. Specifically, we find that stimulation is both phase and state-dependent and that the exact phase-sensitivity of stimulation depends on the current dynamical state. To separate the distinct contributions of phase and state-dependent factors, we contrast the notions of *state morphing* (local phase shifting without state switching) against *state switching* (phase shifting due to global state reorganisation). More importantly, the probabilities of both morphing and switching are phase-, state and region-dependent. Therefore, to correctly predict the effects of stimulation, we need complex mathematical mappings that can account jointly for all these factors (phase, state, region). We show that incorporating dynamical state information in machine learning algorithms improves the prediction of distributed phase-shifting induced by stimulation by up to 40%. Our results suggest that improving the reliability of stimulation may crucially depend on monitoring the transient dynamics of the whole brain, beyond simply evaluating the phase for the activity in the targeted region during stimulation.

## Results

### State morphing versus state switching

Traditionally, the phase-dependency of the phase response to an external perturbation (and, specifically, to a pulse perturbation) has been described in terms of the so-called phase response curve (PRC). The PRC is a transformation function which quantifies how much the phase of an oscillator is shifted in response to any given input depending on the precise time at which the input arrives with respect to the ongoing oscillatory activity, specifically, the phase (Fig 1A; (*27*). Stimuli of identical intensity and duration may advance or delay the oscillation phase of a stimulated oscillator, depending on the phase at which stimulation is applied. These opposing effects are reflected, respectively, in the positive or negative values of the PRC profile for stimulation at different phases.

**Figure 1.**
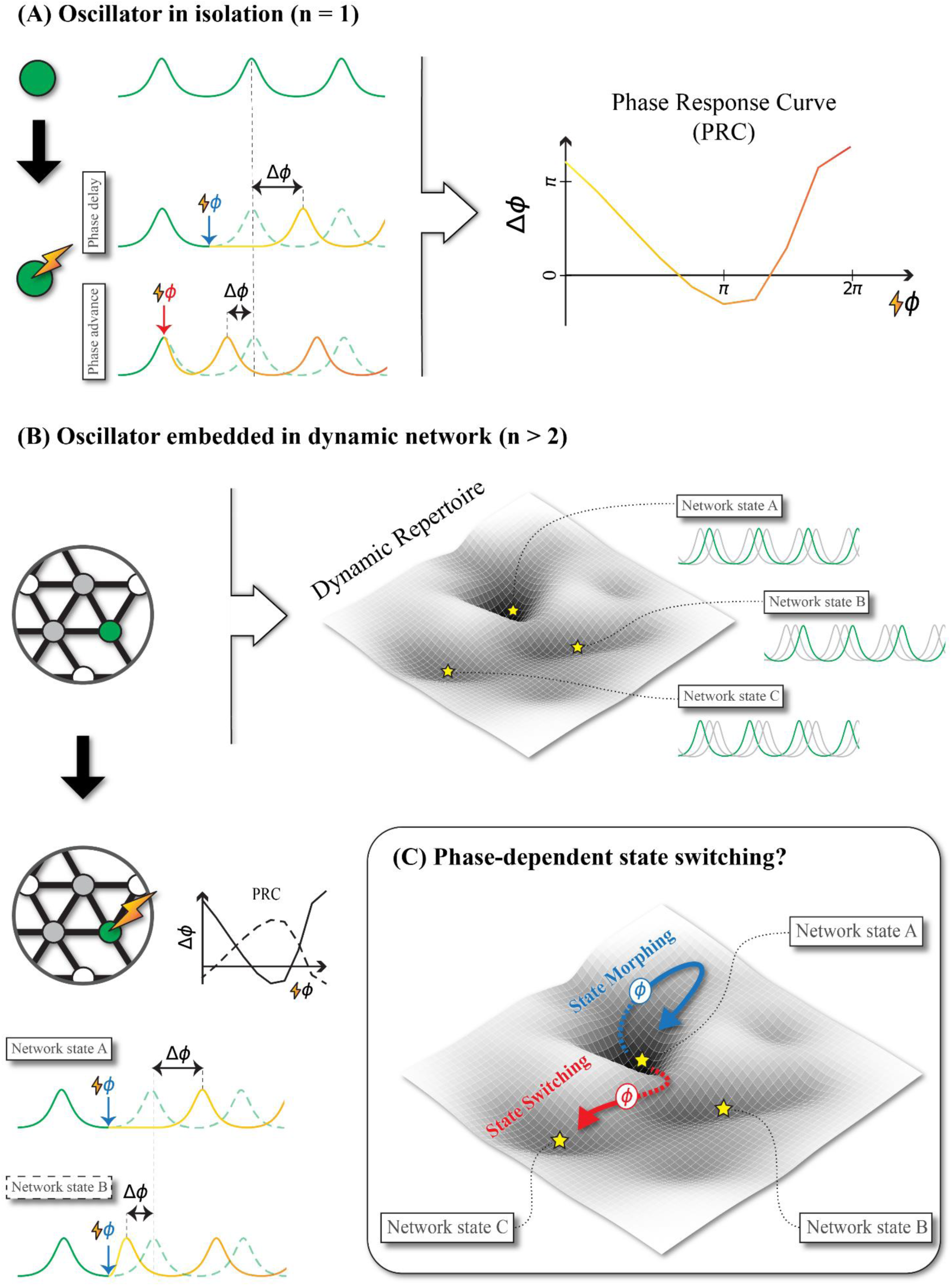
Cartoon representing the effects of single pulse stimulation in an isolated oscillator and in an oscillator embedded in a dynamic network.

The PRC usually obtained from oscillators in isolation, weakly coupled to the stimulation. However, it has been shown to remain pertinent for the case of two weakly coupled oscillators, by accurately predicting alternative phase-locking patterns (*33*) and even capturing the effect of stimulation on their coupling (*32*). Naturally, the question poses itself whether the notion of PRC remains useful in large-scale whole-brain networks, where a multitude of oscillating regions exhibit a rich dynamic repertoire with shifts between distinct global dynamic states (Fig 1B; (*34, 35*)). Beyond the weak coupling limit, we can expect that the dynamics of a local region are modified depending on the global state of the network in which it is embedded. Therefore, we may see diverse “effective PRCs” when we stimulate in different transient states of network dynamics, associated with different patterns of functional connectivity (FC) which characterise the overall oscillatory activity (*36*). Even further, the stimulation may induce a switching of the global dynamic state itself, and, thus, a modification of the PRC (Fig. 1C).

The observed phase shifting after a stimulation could thus be due to a mixture of two different processes. Firstly, oscillation phases and inter-regional phase-differences could be locally adjusted without pushing the system to a different collective state. Thus, FC describing the inter-regional oscillatory coordination would be slightly modified but not radically reconfigured, leading to what we propose calling *state morphing* (*37*). Secondly, the stimulation would change the global dynamical state, leading to *state switching*. Thus, the exogenously induced phase advancing and delaying effects across regions (depicted by dotted arrows in Fig. 1C) would combine with the eventually stronger endogenous reconfigurations of phase differences (solid part of arrows in Fig. 1C) due to changes in state-dependent brain wide FC.

In order to quantify these phenomena, we use a deterministic computational model to remove the disturbance of possible confounding factors usually present in experimental data from methodological or behavioural sources. As we will show, indeed, emergent dynamic complexity alone is already sufficient to enforce complex state morphing and switching behaviours.

The phase-dependent response of an oscillation to a perturbation can be quantified by the Phase Response Curve (PRC). The PRC is a local transformation function that captures how phase of a single pulse stimulation determines the phase shifting post-stimulation relative to the unstimulated phase of the ongoing oscillation. This function is formulated for an oscillator in isolation and the question arises whether this concept holds in a large-scale oscillatory network. Will the PRC remain generic or will it be shaped by the functional connectivity (FC) and give rise to FC-specific “effective” PRCs. Furthermore, will stimulation induce *state morphing* (local phase shifting without state switching), where we could extract meaningful “effective” PRCs that capture only local dynamics, or *state switching* (phase shifting due to global state reorganisation). (A) Left: A single node (green) in isolation oscillates in a consistent pattern of activity (solid green line). If stimulated, the phase of the oscillation (yellow) may be delayed (blue ϕ; + Δϕ) or advanced (red ϕ; - Δϕ) relative to the unstimulated time series (dashed green line) depending on the phase at which the stimulation arrives. Right: The dependence of stimulation phase (x-axis) to phase shifting (y-axis) is very regular and can be captured by the PRC. Shown here, is a biphasic cartoon PRC where a phase delay is represented by negative Δϕ and phase advancement is represented by positive Δϕ. (B) Top: An oscillator (green) embedded in a dynamic network (n > 2; grey and white nodes). Only three nodes of the network are shown for clarity (grey nodes, grey lines). The surface to the right represents the dynamic repertoire, a low dimensional representation of preferred functional configurations among the nodes. The deeper/ darker the point on the surface is the more probable the corresponding functional configuration is. The most probable functional configurations (yellow stars) correspond to a leading position for the green node (Network state C), a lagging position for the green node (Network state B), and an intermediate position for the green node (Network state A). Bottom: The “effective” PRC may look different if the green node is stimulated in Network state B (dashed black line) and in Network state A (solid black line). For example, stimulating at the trough of an oscillation (blue ϕ) may lead to a bigger phase delay in Network state A than in Network state B. (C) Stimulating in a dynamic network state may induce initial deviation from the original functional configuration (dashed blue and red lines) and a subsequent shaping by the network (solid blue and red lines). If Network state A is stimulated at the trough (blue ϕ) *state morphing* may be induced. If Network state A is stimulated at the peak (red ϕ) state switching may be induced.

### Computational model

In order to generate an oscillatory whole-brain model with rich dynamics we combine a weighted, and directed structural connectivity derived from Macaque tractography (Fig. 2A) with a neural mass consisting of two connected mean field models representing excitatory (E) and inhibitory (I) populations (Fig. 2B).

**Figure 2.**
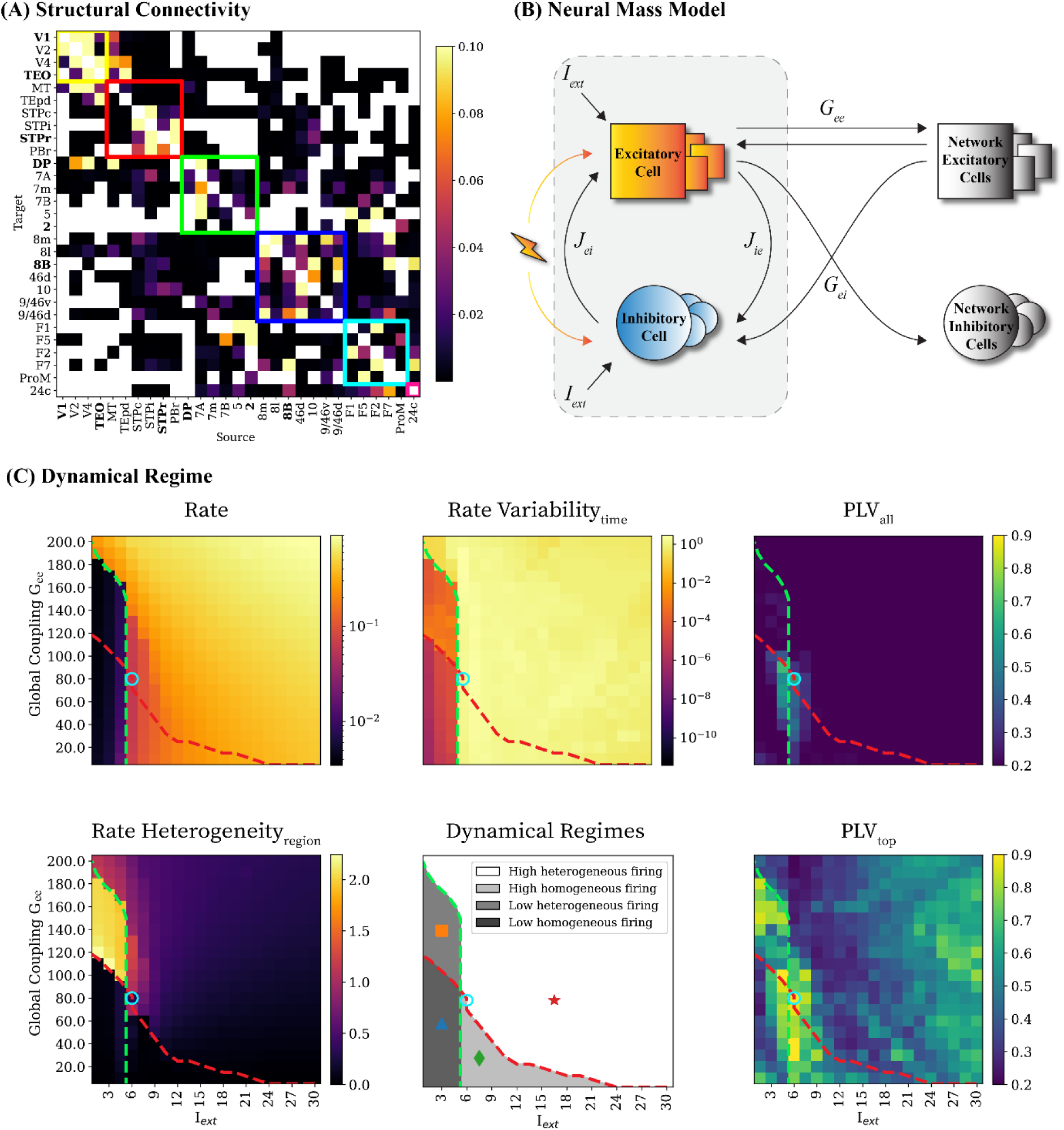
Choosing the dynamical working point of the model.

We choose to use a structural connectome matrix derived from (*38*) because of various reasons. Firstly, it is derived from tracer injections rather than MRI tractography more often used in virtual brain models (*39*). The different reconstruction technique can obtain a connectome matrix which is directed, asymmetric and denser, unlike MRI-diffusion-based matrices, and both these features are expected to boost dynamic complexity, through symmetry breaking (*40*). Secondly, this edge-complete connectome provides ground-truth information about the connectivity between a moderate number of regions, accessible to computationally intensive *in silico* experiments.

To generate plausible oscillatory dynamics within each local region, we connect the E and I populations in a PING-inspired configuration. Furthermore, we use an exactly reduced model for each population so that population level dynamics can be realistically linked with microscopic spiking and synaptic parameters (*42*). Similar to the PING configuration, which is a common canonical description of cortical oscillations in the gamma range (*41*), our neural mass model gives rise to a biphasic PRC (*26, 42*), analogous to the one of Fig. 1A, meaning that stimulation can induce both phase advancements or phase delays when stimulation is applied to both excitatory and inhibitory populations (as occurs for general, cell-type aspecific empirical stimulation).

Then, we explore the effects of global excitatory-to-excitatory coupling Gee and the local background excitatory drive Iext (same for all regions) on the overall network dynamics. In this way, the connectome matrix sets the relative strengths of different inter-regional excitatory connections, but not their absolute strengths (scaled by the parameter Gee). Inter regional connections are purely excitatory but target both the remote E and I populations, with a fixed ratio between Gee and Gei conductance (see *Material and Methods;* Fig. 2B). Together, the two parameters Gee and Iext determine the regime of collective dynamics, which we characterised using various metrics in Fig. 2C and Fig S1.

To explore how state morphing and state switching is shaped by the local phase dynamics and global network states we turn to a deterministic computational model that can generate oscillations. We model each region as a neural mass consisting of an excitatory (E) and an inhibitory (I) population connected in a PING-inspired regime and set the connections between the neural masses with a weighted and directed structural connectivity (n = 28). In order to find a dynamic regime of this model that gives rise to distinct network states we explore the effect of two parameters: The global excitatory-to-excitatory coupling Gee and the local background excitatory drive Iext. We find four distinct dynamical regimes and choose a working point (WP) at the cusp between them which also has a high Phase Locking Value between all regions (PLVall) and a high PLVtop, corresponding to the average of the upper quartile of PLV values (Gee = 80.0; Iext = 6.0). (A) Structural connectivity based on anatomical retrograde tracing data from the Macaque monkey (*38*). Interareal connectivity is marked by squares (yellow: visual area, red: motor area, green: parietal area, dark blue: prefrontal area, light blue: frontal area, pink: limbic area). The names of stimulated regions are written in bold. (B) Neural Mass Model in a PING-inspired configuration. The neural mass has an interconnected excitatory and an inhibitory population. The excitatory population receives a local drive Iext, and projects its firing rate to other excitatory and inhibitory populations scaled by Gee and Gei, respectively. Conversely, it receives input from other excitatory populations to its excitatory and inhibitory populations. Gei is automatically scaled based on Gee. Stimulation is applied to both populations concurrently (see Material and Methods for more details). (C) Dynamical properties of the resting-state network activity for local drive Iext and global excitatory-to-excitatory scaling Gee: Rate, rate variability across time, rate heterogeneity across regions, PLV, and PLVtop. The red dashed line divides regimes whose activity is homogeneous across regions (below) from regimes whose activity is heterogeneous across regions (above). The green dashed line divides regimes who are highly active (right) from regimes who are not as active (left). Symbols mark examples of the four categorical dynamical regimes: blue triangle (homogeneously low firing rate; dark grey), orange square (homogeneously high firing rate; light grey,), green rhombus (heterogeneously low firing rate; intermediate grey), and red star (heterogeneously high firing rate; white). See S1 for example time series for each regime. The blue circle marks the chosen WP (Gee = 80.0; Iext = 6.0).

Simulating time-series in a resting-state (i.e., with constant background inputs and without stimulation), we analyse the firing rate across time and regions. The average firing rate (Fig. 2C, top left) quantifies the overall activity of the network. The rate variability across time corresponds to the firing fluctuations over time (normalised and averaged over regions) as shown in Fig 2C (top, middle), which reflects the oscillation amplitudes. The rate heterogeneity of time-averaged rate across regions is shown in Fig. 2C (bottom, left), with high values for regimes where only some regions are strongly active while others are quiescent. Finally, since the number of strongly activated regions may change over time, we also compute the temporal variability of the spatial rate heterogeneity (S1, top right). See *Materials and Methods* for precise definitions of these four collective dynamics descriptive features.

Inspecting the parameter dependence of these features, we find four qualitatively distinct dynamical regimes, indicated by different symbols and colours in the summary phase diagram in Fig. 2C (bottom, middle), and separated by two crossing dashed lines. The regimes under the red diagonal line display firing rates that are homogeneous for all regions, whereas above it spatial rate heterogeneities start appearing for high values of Gee and Iext. The regimes left of the green vertical line with lower values of Iext have low firing rates as opposed to the right of the vertical line where regimes display higher firing rates. Therefore, we have the following four distinct regimes: with homogeneously low (dark grey, triangle) or high (light grey, square) firing rates; and with heterogeneously low (intermediate grey, rhombus) or high (white, star) firing rates. See S1 for example time-series for each regime from the Working Points (WP) indicated by each symbol.

Besides the oscillatory firing rates, we also characterise the synchronicity of oscillatory dynamics for the same model parameters by calculating the inter-regional Phase Locking Value (PLV), which captures the existence of temporally stable phase differences between pairs of regions (*43*). The PLV averaged over all pairs of regions is generally very low, as indicated by dark blue colour in Fig. 2C (top right). Interestingly, however, in proximity of the crossing of the red and green dashed lines defining the transitions between firing-rate based regimes, the average PLV peaks (reaching ∼0.75). This means that the critical zone in proximity of the convergence between regime boundaries is associated with enhanced global phase-locking. In the following, we will perform stimulation experiments at this WP at the cusp between regimes (Gee = 80.0; Iext = 6.0), whose robust and high global synchrony could facilitate (at least in principle) the detection of stable and reliable phase-dependency effects.

Outside of this line crossing zone, global synchrony is weaker but specific pairs of regions can still be highly phase locked. This weaker type of synchrony can be detected by calculating the average PLV of the 25% of links with the strongest pairwise PLV (Fig. 2C, bottom right). This analysis reveals that smaller subsets of synchronised regions can exist even far away from the critical line crossing point. Remarkably, however, even in the critical zone, the average top 25% of most synchronised pairs of regions reaches values of up to ∼0.89, well above the global average. This finding indicates that the degree of phase-locking is heterogeneous and that there could be cores of highly synchronised regions, raising the question of their stability or transiency over time.

### Transient dynamic state extraction

In order to evaluate and predict the effect of stimulation, our underlying assumption is that the transient dynamic network state must be incorporated into the analysis. This requires the quantitative characterisation and extraction of such states from the collective network activity. We decided to rely on FC, which captures the coordination between pairs of regions, rather than region-specific dynamical properties. FC is thus particularly suitable for tracking system’s level dynamical configurations (*36*) and their evolution in time (*44*). Here, because of the oscillatory nature of the considered dynamics, we define FC in terms of oscillatory phase locking (quantified by PLV between all pairs of regions) and inter-regional phase differences quantified by normalised phase lag (e.g., equal to 0 for in-phase locking and to 0.5 for anti phase locking, see *Materials and Methods).* The rationale is to capture not only the presence of synchronisation, but also information about the extent to which regions precede each other, as we detail now.

To measure FC in a dynamic fashion, we use a sliding window approach (window length = 140 timesteps, overlap = 75%) and then extract our FC measures for each (time) window. For each window and each region, we calculate a “*hubness”* measure, which is larger

To detect the transient global network states which the network is exploring over time, we generate resting-state time series for the chosen WP (Gee = 80.0; Iext = 6.0). Then we use a sliding window approach (window length = 140 timesteps, overlap = 75%) to extract transient windows and calculate phase-based FC measures for each window: Phase-locking hubs (regions that are highly phase-locked to many other regions) and Phase lag (the direction of lag between stable hubs). Jointly, they select three largely overlapping clusters (Ψ1, Ψ2, Ψ3) with highly dissimilar patterns which give us robust targets to stimulate in different transient network states. (A) Left: Cartoon of hub identity for a threshold of three. The presence of links between nodes represents high phase-locking in a window. The number in the nodes corresponds to the number of such links per node. If a node has three or more links it is marked in red. Otherwise, it is marked in white. Right: Hub identity matrix across regions and windows from resting-state time series. X-axis is ordered and delineated with a black vertical line according to five hub states obtained with AgglomerativeClustering.

Hub presence is marked in red and absence in white. Stimulated regions are labelled. (B) Left: Cartoon of lag between region 1 and 2. To align the time-series of region 2 (grey) with the time-series of region 1 (black) it needs to be shifted by lag d (dashed pink line). Right: Normalised lag matrix across regions and windows from resting-state time-series. X-axis is ordered and delineated with a pink vertical line according to four lag states obtained with AgglomerativeClustering. (C) Left: Confusion matrix of hub and lag states. FC states are based on overlapping hub and lag states: Ψ1 (blue), Ψ2 (orange) and Ψ3 (green). Other windows in grey are termed “Other”. Right: Non-supervised, non-linear dimensionality reduction method of concatenated hub identity matrix and lag matrix show three well-defined clusters, which are well captured by the chosen FC states. The five most typical windows for each FC state are marked with a star in the corresponding colour (blue, orange, green). (D) The prototypical FC state with respect to hub probability and lag. Regions that will later be stimulated are marked in bold.

(smaller) for strongly phase-locked regions with many (few) other regions (Fig. 3A, left). We call regions, whose hubness exceeds a chosen threshold, *phase-locking hubs*. Since the PLV matrices change over time (i.e., across windows), we stress that hubness and hub identity are dynamic properties: different regions can be labelled as phase-locking hubs at different times. This is shown in Fig. 3A (right), where red bars denote regions labelled as hubs in a given window. The windows have been reordered according to an unsupervised clustering (in k = 5 cluster) of the time-dependent sets of hubs (see *Materials and Methods* and Fig. S2 for details of hubness analyses), yielding five groups over all considered windows.

**Figure 3.**
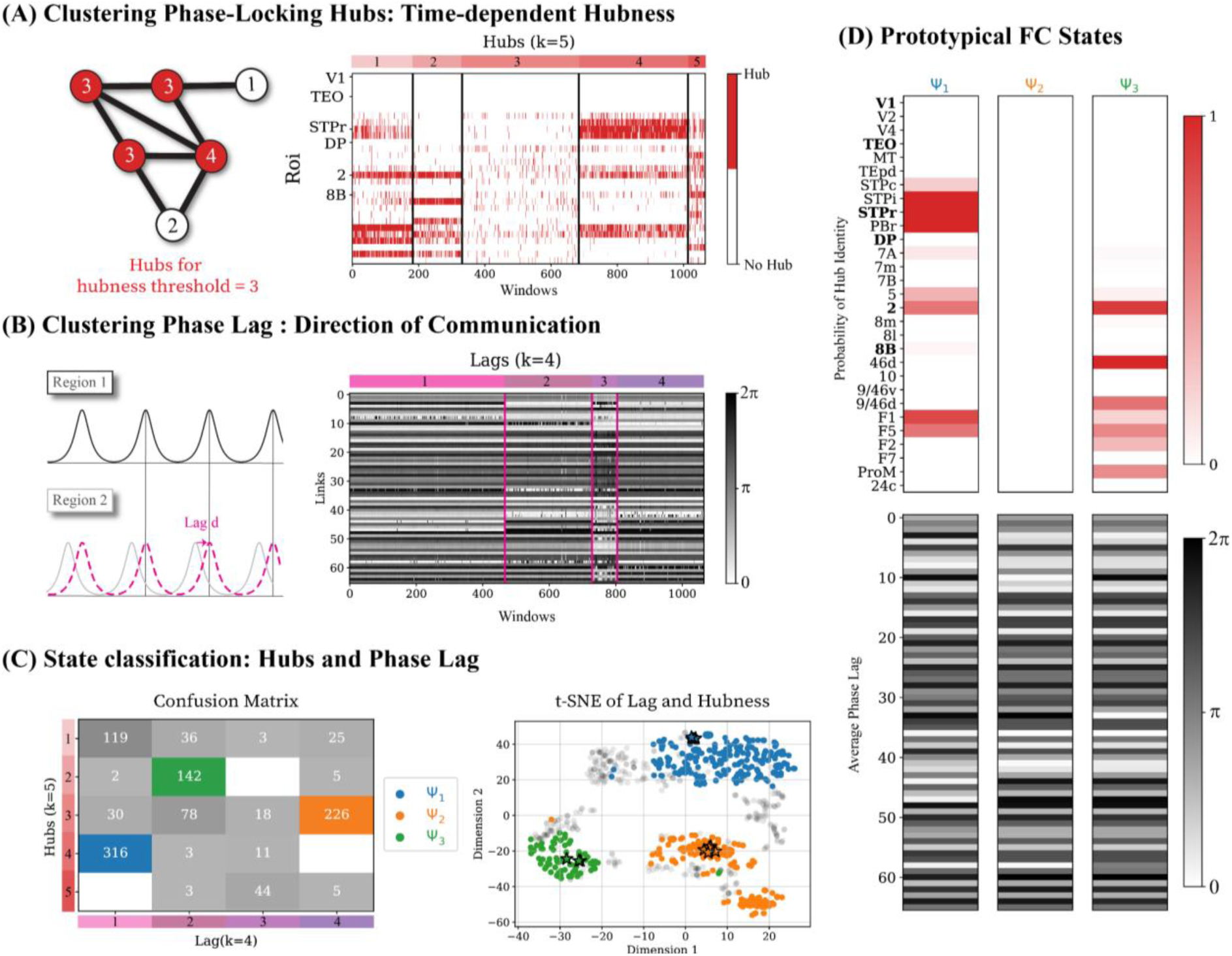
Hub and lag jointly characterise well-defined FC states.

The normalised phase lag (or, in short, lag; cf. Fig. SM3.2) gives a proxy for the direction of communication based on cross-correlation (*45*). Once again, lags are dynamic and the same FC link with strong phase-locking can exhibit different preferred phase differences across windows (Fig. 3B). Vectors of phase lags in Fig. 3B have been reordered in four groups, once again based on unsupervised clustering.

Interestingly, these groups extracted from hubness on the one hand and lag on the other largely overlap as shown by the confusion matrix in Fig 3C (left). This identifies the presence of at least three distinct clusters with well-defined sets of hubs and inter-regional phase relations. These clusters are also easily visible as disconnected clouds (each dot representing a window) in a joint dimensionally-reduced representation of hub sets and phase lags (Fig 3C, right), obtained by t-distributed stochastic neighbour embedding (t-SNE; (*46*)). We refer to these three clusters as states Ψ1 (blue), Ψ2 (orange) and Ψ3 (green). Fig. 3D shows a graphical representation of their centroids, corresponding to stereotypical patterns of PLV-based hubs and lags. Note that these states are not the only patterns exhibited by our dynamic system and we group all remaining patterns under a generic fourth group ‘None’, beside the well-defined states Ψ1, Ψ2 and Ψ3.

Through this double-clustering procedure, every window is attributed to a state label. Inspecting the sequence of state labels over time, we find that states are temporally stable: at each time, it is more probable to remain in the same state in the next window, than to switch to another state (cf. S3); in fact, transitions between states occur via windows from the ‘None’ group. This temporal stability allows us to isolate time-epochs in which the system consistently visits a given state, such that the application of stimulation in such epochs enables the study of the state-dependency of its effects.

### Simulated phase shifting effects depend on stimulation phase, FC-state, and region

In order to examine the state-dependent effects of stimulation we simulate a single pulse stimulation applied within windows corresponding to one of the FC states (Ψ1, Ψ2 or Ψ3). The spread of the state clouds indicates some variability across windows within each state (Fig 3C, right; see *Discussion* and last section in *Results)*, therefore we constrain our study to highly representative windows for each FC state to capture state-specific responses. We thus select 5 typical windows that are most similar to the centroid of each state cloud, while being far from other state clouds (see stars in Fig 3C, right). Within those selected windows, we perform multiple stimulation experiments, applying individual stimulation pulses at different phases in each simulation (see *Material and Methods* for details on stimulation simulation).

We set up a stimulation experiment to explore the role of stimulation phase (0 - 2π, in 10 steps), stimulation FC state (Ψ1, Ψ2, Ψ3), and stimulation region (V1, TEO, STPr, DP, 2, 8B), concurrently. Each dimension can strongly affect the phase-shifting response post-stimulation as we can see from the phase shifting portraits (Δϕ). If we only change one parameter such as phase (A vs B), FC state (A vs C), or region (A vs D) the phase shifting changes completely. However, the changes may be induced by differences in state switching for which we are not controlling yet. Each phase shifting portrait is calculated for each parameter combination of stimulated phase, FC state, and region by calculating the angle of the complex average across five subsequent cycles in the five most typical windows equalling 25 samples. Lightening indicates a stimulated region. Δϕ is wrapped in a range between - 0.5π and + 0.5π. (A) Phase shifting portrait for stimulating region 2 at phase 0.4π in FC state Ψ1. (B) Phase shifting portrait as in (A), but for phase 1.2π. (C) Phase shifting portrait as in (A), but for FC state Ψ2. (D) Phase shifting portrait as in (A), but for region V1. Changing just one parameter drastically changes the stimulation response of the network.

The numerous dimensions explored here (phase, FC state, regions) increase the computational cost of numerical simulation exponentially. Concretely, we chose to explore 10 different phase bins and three different FC states with 25 samples (five different oscillatory period cycles in each of the five different typical windows) for each combination such as to average over multiple instances to quantitatively capture the variability of stimulation effects; this choice yields 750 simulations for each stimulated region. An exhaustive study of all 28 regions would thus require running 21k individual simulations of the dynamics of a whole virtual brain modulated by stimulation. These numbers put a limit on the practical feasibility of exhaustively investigating all combinations.

Therefore, we focus on a subset of six regions from various cortical areas and different levels of phase-locking within the network: TEO in the visual area, STPr in the motor area, DP in the parietal area, and 8B in the prefrontal area (see bold names in Fig 2A). Furthermore, we also added the region with the fewest hubs (V1) and the region with the most hubs (2) thereby covering a variety of regions. Investigating stimulation effects over this subset of regions still required running over 4k *in silico* stimulation experiments, which is a more realistic, yet daunting, endeavour.

To capture the effect of each stimulation pulse we simulate two sets of (multivariate) time-series of whole-brain activity in parallel: the unstimulated and the stimulated time-series that only differ in terms of whether stimulation was applied at a specific phase, region, and FC state. Because our model is deterministic and the parallel simulations start from the same initial conditions, we can be sure that the observed deviations are exclusively due to the stimulation. Furthermore, this determinism allows us to anticipate the phase at which we apply the stimulation with complete precision (an advantage compared to empirical experiments).

To calculate the phase shifting effect of the stimulation we subtract the phase of unstimulated time-series from the phase of the stimulated time-series. We stress that we evaluate the effects after averaging over 25 repetitions for different oscillation cycles, but with the same initial conditions in terms of region, state and phase where the stimulation is applied (see *Materials and Methods*). The phase response goes through a transient shifting behaviour initially at ∼100 timesteps and stabilises after ∼400 timesteps (see Fig 4A). In line with previous studies (*18, 30, 47, 48*), the average phase shifting effect of stimulation is also highly non-local (i.e. regions far away the stimulated one can be affected, cf. (*49*)). Furthermore, it is heterogeneous across affected regions meaning that there is a structured network-level response (see Fig 4A).

**Figure 4.**
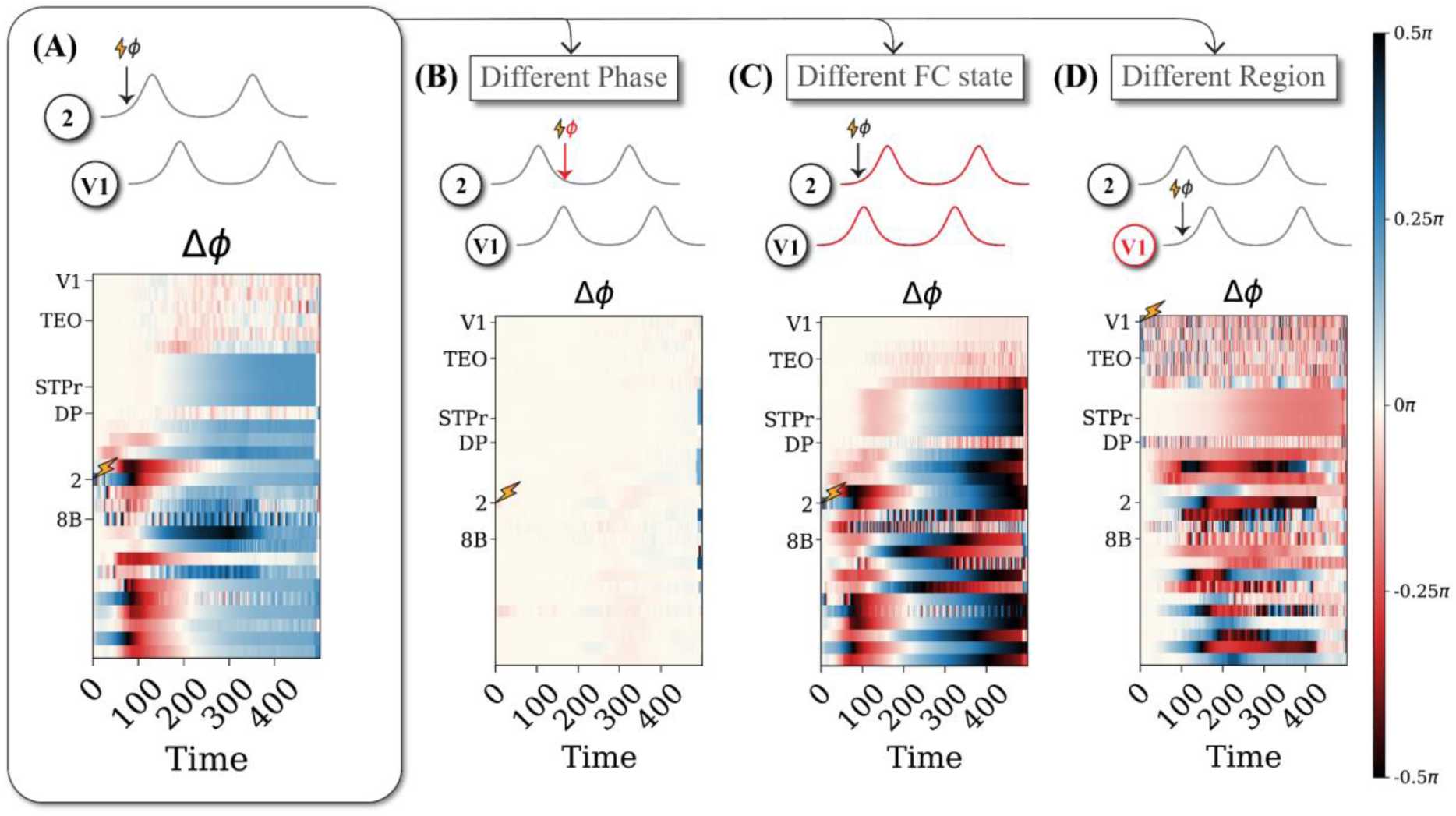
Phase shifting portraits (Δϕ) differ strongly across each dimension of interest (phase, FC state, region).

In line with the concept of the PRC, stimulating at a different phase with otherwise identical parameters can lead to strikingly different patterns of phase shifting across regions (compare Fig 4A and B). While there is a strong network-response in 4A for a phase of 0.4π the response is flat in 4B when stimulating at phase 1.2π for exactly the same region and same initial condition within the same FC state. This confirms the presence of phase-dependent effects in our simulation experiment.

Most importantly, we also see strongly divergent patterns when we stimulate in a different FC state (compare Fig 4A and C), which is a key prediction of our main hypothesis. While both phase shifting time-series display similar initial transients at ∼100 timesteps, the phase shifting trajectories diverge at a later stage. Indeed, after equilibration at ∼400 timesteps, there is a global phase advancement for Ψ1 in 4A, whereas there is a heterogeneously structured phase delay for Ψ2 in 4C.

Lastly, stimulation effects also depend on the region of application as network-wide phase-shifting profiles change when different regions are stimulated (Fig 4A,D). For example, an initial transient response emerges around ∼250 timesteps and not ∼100 timesteps when stimulating region V1 (Fig 4D) and settles as a global structured phase delay and not an advance after 400 timesteps. Some regions such as 8B, DP, and STPr were extremely rigid in response to stimulation for all FC states and phases (S4), whereas other regions were sensitive during specific FC states (V1) or during specific phases (2). Region TEO has some weak stimulation effects while regions such as 8l show phase-independent effects in Ψ2 or regions such as V1 demonstrate both phase and FC-state dependence. These results emphasise that the stimulation responses in a complex oscillatory network are multifactorial and cannot be explained by a single factor.

### Disentangling state switching from state morphing

A possible explanation underlying some of the heterogeneous phase shifting profiles shown in Figure 4 could be that stimulation in different conditions may induce different phenomena. Indeed phase-shifting due to induced state morphing (Fig. 1B) may be more moderate and more local, while phase-shifting due to induced state switching (Fig. 1C) is expected to be more radical and distributed across the whole network. We thus track the state labels for the parallel stimulated and unstimulated time-series in windows following the stimulation to properly discriminate between the state morphing and switching cases. This is possible in our in silico experiments by means of a trained machine-learning classifier: we rely on the state labels obtained via the unsupervised clustering of resting state (unstimulated) dynamic FC matrices (cf. Fig. 3) and use them to generalise the classification to previously those for the stimulated times series (with 3 consecutive windows). The classifier allows us to assign a state to which the transiently observed PLV and lag configuration most likely belongs

The state transition matrices for the parameter combinations in Figure 4 show that differences in state switching may be underlying the differences in phase shifting portraits. Furthermore, meaningful PRCs may not be extracted from trials where state switching has taken place as it would be capturing a mix of global and local dynamics. Therefore, we split trials into *state morphing* where we calculate the “effective” state-dependent PRCs and *state switching* where we calculate the probability of stimulation-induced state switching. We find distinct PRCs for the various FC states and a phase-and state-dependence of state switching. (A) State transition matrices for the same parameter combinations as in Figure 4. The darker the colour the higher the probability is. (B) Diagram distinguishing state morphing from state switching. State morphing is defined as samples that follow the stimulation trajectory on the diagonal, i.e., remaining in Ψ1 (blue), Ψ2 (orange), Ψ3 (green). State switching is defined as samples that follow a stimulation trajectory off the diagonal. (C) PRC calculated based on samples that are exclusively state morphing. Vertical lines corresponding to FC state colours show the standard deviation. (D) Probability difference between stimulated and unstimulated state switching. Probability of state switching in the non-stimulated state switching trials are subtracted from the probability of state switching in the stimulated state switching trials to quantify the effect of stimulation on state switching. For positive values the stimulation induces state switching, whereas for negative values the stimulation prevents state switching. Dots in the corresponding state colour mark the stimulated phases.

to (see *Materials and Methods*). Stimulated trials are then sorted into two groups: trials starting and ending in the same state (phase shifting effects corresponding to state morphing) and trials with a different end state compared to the start state (hence state switching). In this way, the two effects can be disentangled despite the presence of uncertainty in classification. Recall that

this enforced discretization with three FC states is an approximation that neglects the variability within each FC state, which can result in high variability across stimulation effects with the same conditions on phase, state, and region (cf. Fig. S4). We will discuss later this “limitation” of our description (see also *Discussion*), but we focus on the distinction between morphing and switching in the following.

Considering state switching, we consider the same stimulation experiments as in Fig. 4 and quantify the probability of stimulation-induced state switching, rather than a generic mixed-origin phase-shifting. We find that state switching occurs consistently, for the example in Fig. 4A from state Ψ1 to Ψ2 in 96% of the cases (Fig. 5A, first column) and, for the example in Fig. 4C, from state Ψ3 to Ψ2 in 100% of the cases (Fig. 5A, third column). However, state switching does not occur for the examples shown in 4B and 4D (Fig. 5A, second and fourth columns, respectively). This confirms that the probability of state switching also depends on phase, state, and region as in Fig 4.

**Figure 5.**
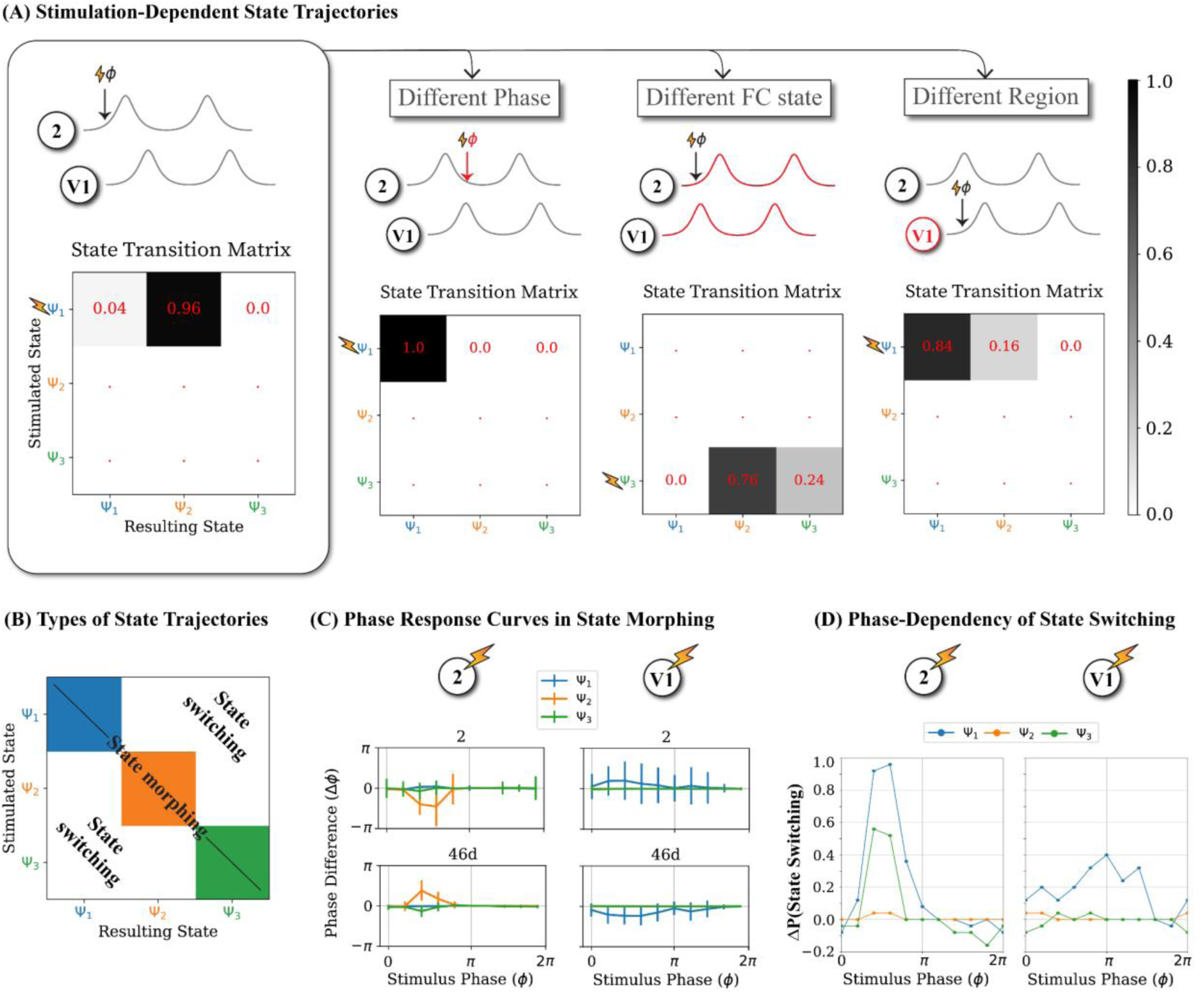
Stimulation can induce both state morphing and state switching.

### State morphing is shaped by the dynamic FC-state

Now focusing on stimulated trials without state switching, we study the weaker phase shifting effects associated with state morphing. To discard transient effects, we measure stimulation-induced phase shifting in a late post-stimulation window (360 - 500 timesteps after stimulation) and compile these measurements into PRCs describing their phase-dependency, state-by-state. Even after excluding all trials with state switching, the extracted PRCs are strongly affected by FC states (Fig 5C). For instance, when region 2 is stimulated in FC state Ψ1 and Ψ3 its phase is unaffected; however, when stimulating it in FC state Ψ2, we observe consistent phase delays for stimulation phases around ∼π/2 (Fig 5C, centre left). Expectedly, the PRCs depends on both the stimulated region and the affected region as shown for state Ψ2 with phase advancements versus delays (Fig 5C). While the PRCs for STPr, 8B, DP, and 2 show very consistent responses across samples, those for TEO and V1 are dominated by variability of larger magnitude than the average phase shift (see S6) due to large single trial variability (Fig S5). This observation indicates once again (cf. Fig. S5) that description of transient activity can be refined by going beyond the defined discrete states.

### State switching is dependent on the phase of stimulation

Following Fig. 5A with dependency of state switching upon phase, FC state, and region, we further represent this information in terms of curves for the phase-dependent probability difference of switching (Fig. 5D), which is more directly comparable with PRCs associated with (state-dependent) state morphing (Fig. 5C).

Interestingly, regions with a high probability of stimulation-induced state switching such as region 2 (and V1) also show a strong phase and FC state dependence in their state switching probabilities (See Fig. 5D, left; S7). For example, in region 2 states Ψ1 and Ψ3 are very likely to switch to another state when the stimulation arrives at phase ∼π/2, whereas Ψ2 is much more robust and its switching deviation is uniformly low for all phases and stimulation even seems to supress natural state switching (for negative values). In regions such as 8B (or DP, TEO, and STPr) for which the propensity of stimulation-induced state switching is lower, there was also no strong phase-dependence (See Fig 5D, right; S7). Overall, strong state switching deviations can be induced by applying stimulation in specific combinations of phase windows and regions, as previously observed by other studies (*32, 50*).

Clearly, FC states are an important dimension to consider when attempting to understand the effect of stimulation in a complex oscillatory network. The results in Fig. 5C-D show the limits of the traditional PRC concept and how the effects of stimulation transcend it. On the other hand, it may quickly become impractical to build a dictionary of all possible state dependent PRCs in large networks. In order to understand and predict stimulation effects, we thus need to introduce more straightforward strategies, explicitly accounting for the existence of complex dependencies for multiple factors simultaneously.

### State-aware information is essential for predicting stimulation-dependent phase shifting

To condense the rich multidimensional information contained in our simulated stimulation experiments and make practical predictions on the resulting effects, we rely on machine-learning tools. Specifically, we train a Random Forests Regression algorithm (see *Materials and Methods*) to predict the stimulation-induced phase shifts using cross-validation based on a flexible combination of input variables, which describe the network with various levels of granularity.

We compare the prediction capabilities of input variables that are “state-ignorant” versus “state-aware” (see purple and pink boxes in Fig 6A). The former describes local features like the phase of the stimulated region (w.r.t. its own oscillation) at the time when stimulation is applied. We also consider the outgoing structural connectivity of the stimulated regions. The rest of the variables describe the global FC state at the time of stimulation in various levels of detail like the FC state identity, PLV and lag matrices (or summary metrics derived from them).

**Figure 6.**
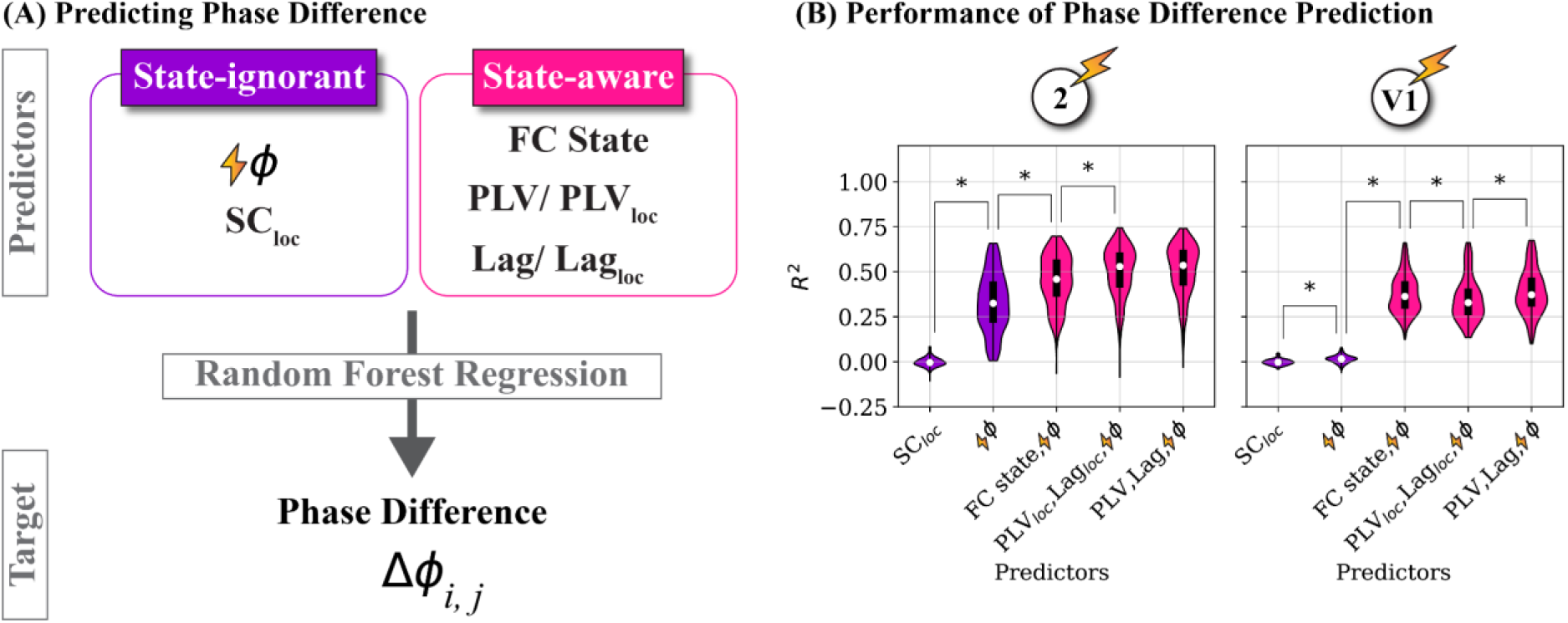
State-aware predictions can significantly improve the prediction of stimulation effects.

To quantify the relative importance of information about the network dynamic state and to predict the effect of stimulation across FC states we train a Random Forest Regression (RFR) algorithm to predict the phase-shifting with respect to the unstimulated time series in the most posterior window of the stimulation trials. (A) Top: Predictors for the regression can contain local information (state-ignorant; purple) such as stimulation phase and outgoing structural connectivity of the stimulated region (SCloc) or may contain information about the transient network state (state-aware; pink) such as FC states, the full PLV and lag matrix, or the PLV and lag links with regions that the stimulated region was structurally projecting to (PLVloc, Lagloc). Bottom: These predictors can be flexibly concatenated to predict the stimulation induced phase shifting. (B) Prediction performance of stimulation-induced phase-shifting.

We used RFR with a depth of 5 and used 10-fold cross-validation (training size = 0.7) to quantify the performance of each affected region independently. Violin plots reflect the test performance of 10 CV folds for 28 affected regions (280 points in total). Including state aware predictions significantly improved the prediction. In each violin plot the white dot marks the median and the thick black line marks the inner quartiles. Significance was tested with the Mann-Whitney U test and significance levels are indicated as follows: not drawn for p ≥ 0.05; * for p < 0.05. Furthermore, only significance tests for adjacent distributions are drawn (See S8 for full set of comparisons). Bonferroni correction for the full set of multiple comparisons was applied.

The predictive power is significantly improved by the state-aware variables (pink) with respect to the state-ignorant variables (purple), as shown by the comparison of prediction performances in Figure 6B (for stimulation in two example regions, V1 and 2, see S8 for other regions). For the state-ignorant variables, stimulation phase outperformed the outgoing structural connectivity of the stimulation region SCloc for all stimulated regions (except DP). Furthermore, when including state-aware features performance increases up to ∼ 35% for stimulating region 2 and region V1. Note that the relative performance of state-ignorant and state-aware predictors heavily depends on the stimulated region, with strongest improvements for V1, 2, 8B, and STPr (∼ 20 % to ∼ 40 %), compared to moderate ones for regions DB and TEO, in a range between ∼ 7 % to ∼ 14 % (Fig S8). Specifically, we considered three possible alternative state-aware inputs. First, we used just the discrete label of the FC state (Ψ1, … Ψ3) attributed to the time epoch in which stimulation is applied. Even such a rough indication of the whole network dynamics is already enough to significantly improve the performance as compared to state-ignorant prediction. A further improvement is obtained by using: second, the description of the local neighbourhood of the stimulated node in the FC network,, characterised by the PLV values and the phase lags of this node *i* with all the other regions structurally connected to it (i.e. the *i*-th rows of the time-epoch specific PLV and Lag matrixes); or, third, the entire time-epoch specific PLV and Lag matrices (including as well PLV values and lags between pairs of regions both remote from the stimulated node). Interestingly, the size of improvement from state-ignorant input variables (stimulation phase) to extracted FC states labels and PLV and lag matrices was not systematic across stimulated regions. For example,

there was no significant improvement with full PLV and lag matrices with respect to the discrete FC states in V1, whereas region 2 improved with every more fine-grained state-aware input variable. This suggests that for some regions limited information is lost by discretizing the dynamical repertoire into FC states. Another point is that adding only the structurally connected PLV and Lag values as state aware predictors generally performed as well as using the whole PLV and Lag matrices (with exception of V1). The two choices of input features may end up not being very different, given how dense the SC matrix is (cf. Fig 2A). However, another possibility is that, indeed, accounting for the interactions of the stimulated node with just its local neighbours provides a reduced description sufficient to capture system’s level complexities. In the context of statistical physics, a description in which the local neighbourhood of the stimulated node is parameterized corresponds to a so-called “Bethe Peierls” approximation (*51*). This level of description is already more detailed than a “mean field” approximation in which the global network state is just compressed out into a single label (i.e., FC state identity). It is however less detailed than the “exact” description in which the system’s state is represented in full detail by the complete PLV and Lag matrices. Thus, we could summarise by saying that for predicting the effect of stimulation: a mean-field account of the system’s state is already better than ignoring it; and that a Bethe-Peierls account of the system’s state may already be sufficient for further improvement without the need to go towards an exact description (see *Discussion*).

Lastly, the results shown in Fig 6B report prediction performances irrespective of the state in which the system was when stimulated. It is possible, however, that the classifiers perform better when stimulating the system in certain states rather than others. In other words, the predictability of stimulation effects may be higher or lower depending on the system’s state. We thus evaluate prediction performances for each of the stimulated FC states separately. As shown in Fig. S10, for all stimulation states, differences in performance are not systematic. Therefore, we conclude that the superior performance of state-aware predictors with respect to state-ignorant predictors holds in general and does not depend on state-ignorant classifiers performing poorly in some of the states.

## Discussion

This paper analyses the contributions of global and local dynamics on the long-term effects of single pulse stimulation in a large-scale computational brain model. First, we characterise the complex oscillatory behaviour of the resting-state by detecting transient network activity states in the dynamic repertoire of the model based on flexible patterns of phase-locking between regions. Thereafter, we show that stimulation effects are shaped by these states. Stimulation could induce *state morphing*, meaning shifting phases while remaining in the network state at the time of stimulation, and *state shifting*, meaning shifting phases across the network in alignment with a different network state that deviates from the resting-state trajectory. In both cases stimulation effects strongly depend on stimulation phase, network state, and region. Finally, we demonstrate that including state-aware information during the prediction of stimulation-induced phase shifting improved performance by up to 40% validating the importance of this often-neglected global information in the understanding of stimulation effects.

Our results are in line with previous experimental studies that have shown. independently, that stimulation phase (*15–21*) and global oscillatory states (*4*) modulated by cognitive or emotional tasks (*28, 52*), can affect the efficacy of stimulation. Furthermore, our complex oscillatory model shows that these factors can even display complex nonlinear interdependencies and should thus be jointly considered when evaluating the stimulation responses in the brain.

The non-linearity in our system limits the applicability of linear control theory as it transcends temporal dynamics such as exponential growth, exponential decay, and sinusoidal oscillations which can be accounted for in the framework of network control theory (*53*). Additionally, tools for inferring causality are strongly grounded in linear systems theory and can only be applied if a system behaves linearly (*54*). Nevertheless, we can apply the notion of causality in subspaces of non-linear systems that behave in an approximately linear manner (*55*). The FC states in our study seem to be a sufficiently detailed characterisation of the complexity inherent in our model. Indeed, each state corresponds to a well-defined cluster of similar dynamical patterns, defining a “basin of attraction” where the system’s dynamics can be approximated via standard tools (e.g., the PRCs). At the same time, the definition of a multiplicity of states allows us to discretise alternative local “patches” (with alternative local causal structures) as FC states within the broader high-dimensionality space in which the system’s nonlinear dynamical flow unfolds (Fig 3C, left). In this way, despite the inherent nonlinearity, we can still model causal interdependencies state by state, which improves prediction significantly for all regions, with respect to state-ignorant attempts to anticipate stimulation effects (Fig 6B).

Note that our definition of “state” is unconventional and specific, linked to tracking the transient dynamics of FC (*44*). This may contrast with alternative notions in the literature. In experimental studies, the term *state* has been used more often to refer to states of consciousness such as wakefulness, sleep, or anaesthesia (*56, 57*) or behavioural states such as resting or task state (*58, 59*). The dependency of stimulation effects on dynamical states had also been addressed by previous computational works (e.g., (*49*)), however these previous works primarily explored alternative choices in global dynamical WP, closer or farther from some critical point and did not focus on the temporal dynamics. Here, we define states in a different way, equating them to transient functional configurations with similar FCs. This has the advantage of accounting for fast fluctuations of dynamic organisation that occur even within global brain states (*60, 61*). Framing the ongoing activity in the brain as the stochastic sampling of an internal repertoire of alternative states, quantifiable in terms of dynamic FC analyses, emphasises the existence of a largely neglected source of variability in stimulation effects. As such, it may redefine the way in which future studies could be designed to detect this alternative type of state-dependency which we predict in this study.

Remarkably, such rapid variability of FC states occurs in our model in absence of any synaptic plasticity. This may call for a reinterpretation of the mechanisms behind the stronger efficacy of repeated stimulation. Stimulation studies are preferably conducted with repetitive stimulation as the effect is larger and more persistent, a fact often attributed to plasticity changes (*62, 63*). Plasticity could in principle be added to our modelling. However, even now, our model hints at an alternative mechanism that could explain the superior efficacy of repeated stimulation, without the need of invoking plasticity. Indeed, repeated stimulation could induce cascades of state switching that would force the global system dynamics toward the most temporally stable transient FC states (as it is the case for chains of stochastic transitions converging toward the more attracting configurations). This enhanced stability of FC occurs on a timescale faster than the one of plasticity solely through dynamic self-organisation, which could later be sustained by long-term plasticity changes.

Furthermore, our simulations and theory also suggest that long-lasting effects could also be achieved with single pulse stimulation, provided that the timing of this stimulation is properly chosen, accounting for ongoing local and global dynamics. Our model indeed predicts that the effect of single pulse stimulation will be weak for most combinations. Nevertheless, it may overcome the rigidity of the brain state (*10*), meaning the failure of eliciting effects, at specific “dynamic hotspots” (i.e. a particularly stimulation-sensitive combination of phase and FC state). Detecting such hotspots calls for an online tracking of dynamic FC state prior to stimulation (*32, 64–66*), which may be feasible if states dictionaries are pre-extracted from resting state time-series and simple distinctive features identified (e.g. state-specific hubness of a few reference regions, cf. Fig. 3A and 3D).

In our study, we made the operational assumption that discrete FC states can be defined (Fig. 3), an assumption which led us to the distinction between state morphing and state switching, following terminology first introduced by (*49*). However, whether FC states are discrete or continuous is a long-standing debate in the dynamic FC literature. Many authors have used unsupervised clustering to extract discrete FC states (e.g. (*67, 68*)) and alternative formalisms describing dynamic FC as a smooth reconfiguration process have been proposed (e.g., (*69*)). Here, we show that characterising FC states as discrete states is already enough to improve the prediction of stimulation-induced effects (Fig 6B). However, characterising FC states as continuous by using raw instantaneous measures such as PLV and Lag (instead of discrete FC state labels) can further improve prediction, at least for some regions. The co-existence of epochs in which FC configurations are sufficiently stable to be approximated as discrete states with other epochs in which the FC evolution is faster and visiting atypical configurations is reminiscent of the alternation between metastable “knots” and fast reconfiguration “leaps” previously described in resting state dynamic FC (*69*). In the absence of truly discrete states, the existence of transient stabilisation around a few specific system configurations still justifies distinguishing state morphing and state switching as two extreme types of stimulation effects, although their distinction may not be so distinct and could rather be described as a continuum of possibilities between them.

From a dynamical systems perspective, this graded variation between discrete-like and continuous aspects in FC dynamics is also compatible with expectations from the Structured Flows on Manifolds theory (*70*) or effective free energy landscape models (*71*). If the system’s dynamical flow is trapped in a slow subspace, or, equivalently, in a deep and narrow valley on an effective energy landscape (cf. Fig 1C, Network state A), then the discrete characterization will be sufficient as intra-state variability will not be large and the stimulation response will be consistently shaped by robust endogenous force fields. On the other hand, if the dynamics unrolls in faster subspaces and samples broad and shallow valley on the landscape (cf. Fig 1C, Network state B), then a continuous characterization will be more informative as intra-state variability will be large and the stimulation response can vary widely in a less constrained manner.

Specifically, intra and inter-state variability may not only be reflected in different phase configurations but can also be reflected in concurrent changes in frequency and amplitude profiles. Frequency may be particularly important for the case of rhythmic stimulation, as it can induce frequency-specific state changes (*72, 73*) and cross-frequency effects beyond the targeted frequency band (*48*), as predicted by other models (*49*). In our analysis we do not consider frequency variability and cross-frequency effects. However, our model is not a simple network of Kuramoto-like phase oscillators with prescribed frequency (as in e.g. (*74*)). On the contrary, oscillations are emergent, and their frequency dynamically tuned by the received inputs. As such, our model does, in principle, give rise to complex frequency modulations which are intrinsically coupled as they originate from the collective dynamics of the same nonlinear dynamic system. Future studies could expand our analysis for characterising dynamic cross-frequency coupling, that can be rigorously studied in exactly reduced models (*75*) similar to the ones we use by using analytically estimated population level PRCs (*76*).

Another important aspect that may be considered in stimulation in conjunction with phase is pre-stimulus amplitude. Experimental studies have shown that, besides phase, stimulation effects can depend exclusively on oscillatory power (*77–79*), the interaction of phase and power (*80, 81*) or neither of them (*82*) exhibiting an ambiguity reminiscent of the complexity displayed in our study. Again, this could be studied in our model in the future as it displays intrinsically coupled phase and amplitude fluctuations, unlike simpler networks of phase-oscillators.

Following this line of thought, it is worth noting that extensions of the PRC concept that can account for amplitude variations have already been extensively studied (*83*). Indeed, using mathematical tools that extend the conventional assumptions behind standard PRCs (isolated node, phase-reduced node dynamics…), descriptions of the stimulus response of an oscillating population embedded within a larger network could be potentially derived in a mathematically rigorous manner. As shown by (*26*), it is possible to rigorously define a “global PRC” that quantifies how the phase-shifting effect induced by stimulation propagates from one population to other populations, in a network of phase-locked oscillating populations. Such analytical treatments have only been proposed for toy models made up of only a few regions. However, this approach may quickly become infeasible as soon as the number of coupled populations increases and the coupling between them becomes heterogeneous and irregular, as in typical connectome-based models. Here, we deliberately chose to abandon formal rigour to introduce an operational notion of *effective* PRC instead. Effective PRCs are not analytically evaluated from regional dynamics equations and network couplings. On the contrary, they are empirically measured in simulated stimulation experiments. The advantage of this pragmatic approach is that effective PRCs can be used to describe the phase-dependency of stimulation effects to an embedded node “as if” this node was isolated. The drawback is that these effective PRCs become inherently state-dependent, as they absorb the complexity of collective dynamics. Thus, effective PRCs allow us to temporarily ignore system complexity and predict the phase-shifting effects of a stimulation as if they were conventional PRCs, provided that the correct, state-specific effective PRC is extracted, and the applied stimulation induces only state morphing without state switching. The real macroscopic PRC of each regional node within our model can only be modified by altering local parameters, such as intra-regional E and I coupling strengths or synaptic delays (*26, 76, 84*) which we do not alter in this study. Therefore, the real PRCs do not change by construction. But in practice the effective PRCs do change, because of dynamically changing inter-regional input currents and transient system configurations.

We chose a “well behaved” dynamic WP to increase the probability of falsifying our central hypothesis and finding that a description in terms of just one classic PRC is still valid, without the need of introducing multiple effective PRCs. To do so, we purposefully selected a regime that was maximally synchronous, as to stabilise inter-regional phase locking relations and increase the chances of observing a stable phase-dependence in the effects of stimulation. Ultimately, however, even in such a well-behaved WP, we encountered a tremendous amount of complexity. We did not perform simulations in other WPs, because the numerical experiments were computationally heavy. Regardless, we expect the complexity in other less synchronised regimes to further increase making our results even more pertinent. Another ingredient missing in our model are delays (*85*), which would also increase complexity as they favour symmetry-breaking, out-of-phase locking and multistability leading to a larger number of states in the dynamic repertoire (*26, 50*).

Overall, our model remains very abstract with respect to several details. Firstly, we are modelling stimulation as a positive, instantaneous, and perfectly localised pulse in an attempt to emulate Dirac’s delta in the classic PRC. In practice, stimulation has a temporally heterogeneous shape and may be positive and negative over the time course of the stimulation (*73*). Furthermore, the primary area of stimulation may not be spatially localised, but rather spread out (*86*). These aspects may be taken into account by future studies and easily modelled, for example, using already implemented features of The Virtual Brain (TVB) neuroinformatic platform (*87*), such as its built-in stimulus editor and surface-based neural field simulator. Another limitation is the strong intensity of the stimulation pulse relative to the ongoing resting state oscillation. However, we expect that weaker more realistic stimulation intensities could still be effective when applied in the ’’dynamic hotspots’’, because the stimulation effect (Fig 1C, red and blue dashed line) may be amplified by intrinsic self-organised dynamics (Fig 1C, red and blue solid line). Secondly, we use a very dense and non-human connectome. However, similar dynamic complexity has been replicated with human connectomes of a lower density (*35, 88*). Furthermore, the low density of connectomes derived from MRI-based tractography could be corrected by using effective connectivity approaches to re-weight individual structural connectivity links (*89, 90*), including those potentially missing in the original connectome matrix

In conclusion, this study could serve as a first step towards using computational models and dynamic aware prediction algorithms to improve stimulation protocols. Traditionally, single trial TMS variability is discarded as noise and averaged out. Our study points towards the possibility that this variability could be a signature of complex dynamics and a useful target for optimising the efficacy of stimulation. In the future, spontaneous and stimulation-induced variability could be used as a fitting target for the construction of personalised models of the stimulation effects in specific patients (as already attempted in the modelling of epileptic seizure spreading, cf. (*91*)), with improved parameter identifiability (*92, 93*). Such personalised models of nonlinear brain dynamics could be adopted to design tailored protocols of stimulation by targeting the patient-specific dynamic hotspots and increasing energy efficiency by reducing the number and intensity of the applied pulses. We expect personalised virtual brain models to achieve better performances in designing control schemes for brain states, beyond current proposals based on linear control theory (*94*). Last but not least, the personalised fitting of virtual brain models to spontaneous and stimulated activity could enable us to reverse engineer directly linked physiological parameters of interest. Indeed, in exactly reduced models as the ones we use here to describe regional dynamics, parameters in the macroscopic population activity-level description can be precisely related to microscopic parameters as neural excitability and synaptic conductance (*95*). As stimulation can also be used for revealing potential pathologies (*96*), we can link the altered stimulation responses to maladaptive circuit mechanisms inferred via model fitting.

## Methods

We implemented all the simulations in *The Virtual Brain (TVB;* https://www.thevirtualbrain.org) software, a framework for the simulation of dynamics in large-scale brain networks, constrained by an imposed structural connectome, with customizable choice of local regional dynamics and possibility to apply arbitrary spatiotemporal patterns of stimulation (*87*).

### Structural Connectivity

Due to computational constraints, we selected a small, but dense structural connectivity matrix (28 areas; see Figure 2A). The anatomical connectivity was obtained from 29 macaque monkeys through retrograde tracing and is weighted as well as directed (*38*).

### Neural Mass Model

To model a single brain region, we coupled an inhibitory and excitatory population described by a mean field approach obtained from an exact reduction as described in (*26*). This model is implemented in TVB under the regional dynamics type DumontGutkin. Within each region, an excitatory and an inhibitory population were coupled similar to a Pyramidal Interneuron Network Gamma (PING) configuration with a local drive Iext to the excitatory population (see Figure 2B), which is a canonical description of cortical oscillations in the low gamma frequency range (*26*). Unlike the traditional PING our configuration also features a local drive Iext to the inhibitory population. However, the local dynamics of this PING-inspired configuration remained markedly similar to the local dynamics of the traditional PING configuration as revealed by stimulation of this configuration in isolation.

The excitatory firing rate of each neural mass was coupled with the excitatory and inhibitory populations of all other structurally connected neural masses. Stimulation entered the neural mass as an additive term in the average membrane potential Ve/i similar to the local drive Iext. Individual inter-regional connections strengths were dictated by the structural connectivity matrix scaled by adjustable free parameters. Long-range inter-regional excitatory connections targeted both the excitatory and the inhibitory postsynaptic populations, with distinct scaling factors Gee (for excitatory-to-excitatory coupling) and Gei (for excitatory-to-inhibitory coupling). The latter is unusual for computational models implemented with the TVB software and thus required a specific adaptation of the source code. Such excitatory-to-inhibitory coupling, besides its realism, contributes to make the collective dynamics of the model richer and more complex. In reference to well-documented ratios of excitatory to inhibitory populations in the brain, the input to the inhibitory population is scaled to be five times larger than Gee (*97*). All connections were instantaneous (no delay was used) and all simulations deterministic (no noise). Therefore, all FC variability stems from dynamic complexity rather than from stochasticity of inputs.

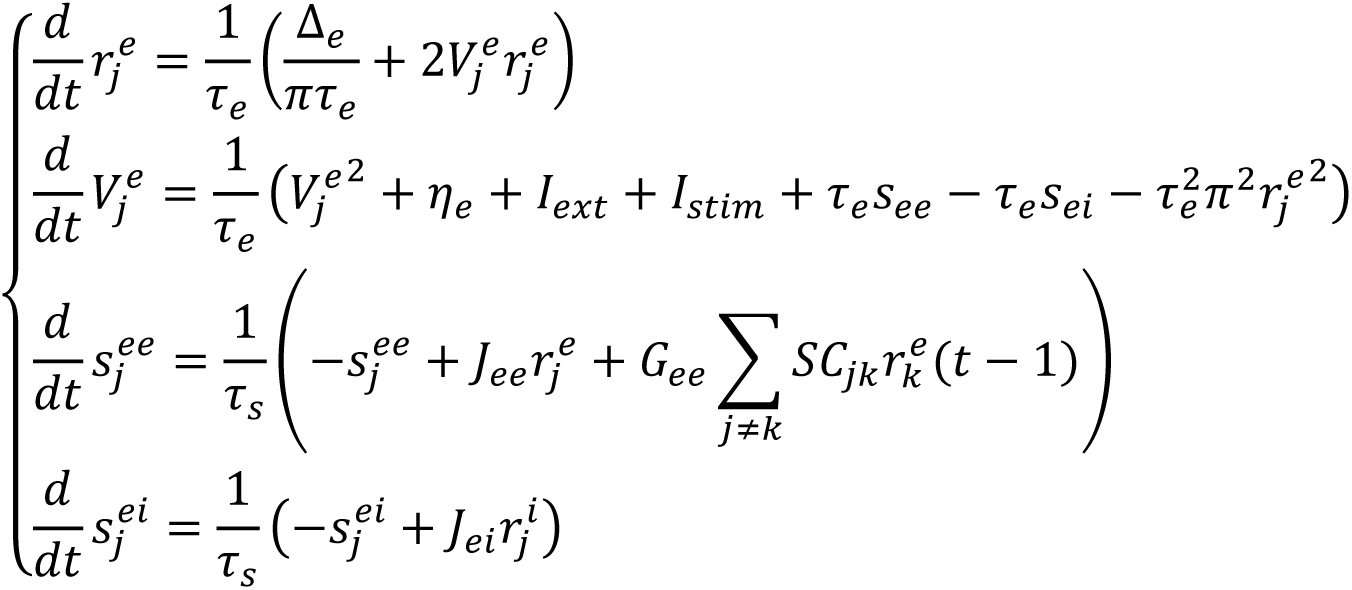

And

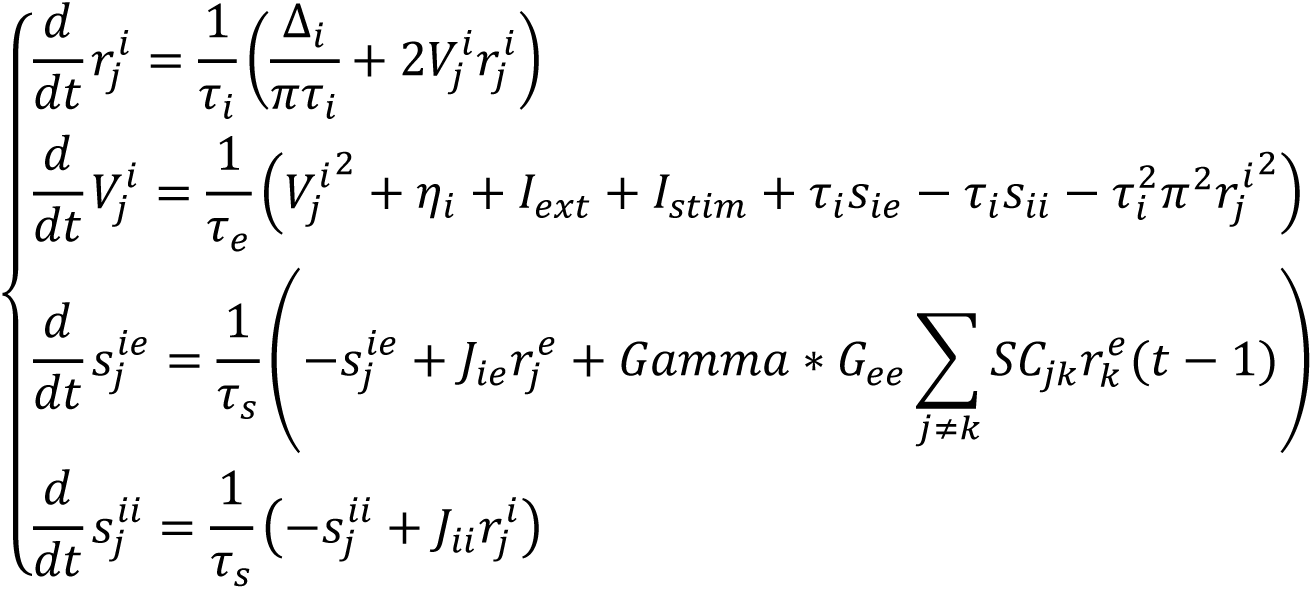

With four state variables per population for region *j*: the firing rate re of the excitatory and ri of the inhibitory population, the average membrane potential Ve of the excitatory and Vi of the inhibitory population, the synaptic currents projecting from excitatory-to-excitatory population (see), excitatory-to-inhibitory population (sie), inhibitory-to-inhibitory population (sii), and inhibitory-to-excitatory population (sei). Other parameters are the spread of the heterogeneous noise distribution of the excitatory population Δe = 1 and the inhibitory population Δi = 1, the mean heterogeneous current to the excitatory population ηe = -5 and to the inhibitory population ηi = -5, the characteristic time constant of the excitatory population τe = 10 and the inhibitory population τi = 10, the synaptic time constant τs = 1, the synaptic weight between the inhibitory population and the excitatory population Jei = 19 and between the excitatory and inhibitory population Jie = 19, the ratio Gei the inhibitory global coupling to Gee the excitatory global coupling Gamma = 5, and the structural connectivity SC as described in the previous section. Finally, the two parameters we explore in search of a WP: the external homogeneous current to the excitatory population Iext and the excitatory global coupling Gee. The equations were adapted from (*26*) and can be consulted for further details how this neural mass was obtained through an exact reduction from spiking models.

### Parameter Exploration: Dynamical Measures

To identify the different regimes of collective dynamics produced by our model, we performed simulations of spontaneous (unstimulated) dynamics systematically varying two free parameters over a prescribed range of values. We explored the global excitatory-to-excitatory coupling Gee versus the local excitatory drive Iext and chose a dynamical regime (Gee = 80.0; Iext = 6.0) at the cusp between clearly defined regimes identified through inspection of various metrics of the generated dynamics of the firing rate in the excitatory population: rate of activity, and its variability across time and between regions; and PLV (Fig 2C). We now describe these metrics in detail.

#### Rate

To quantify the overall level of activity in the network, we calculated *μj* the average rate of activity across time for each region. We then took the median across regions of these regional time-averages (see Figure 2C, top left):

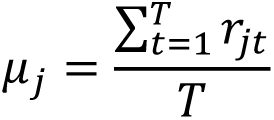

Where *rjt* corresponds to the excitatory rate of region *j* at timestep *t* and *T* corresponds to the total number of timesteps in the timeseries.

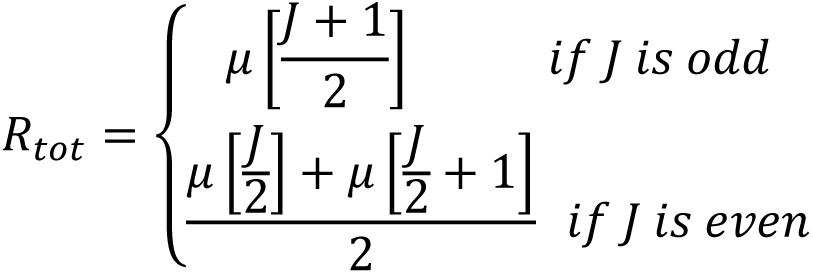

Where *μ* corresponds to the ordered list of average rates of activity for all regions and *J* corresponds to the total number of regions in the dataset. Larger values of Rtot indicate larger overall levels of activity within the network.

#### Rate heterogeneity across regions

To assess whether all regions have similar or different levels of activity, we calculate the Coefficient of Variation (CV) of the region-specific time-averaged rates *μj* (see Figure 2C, bottom left), i.e., the ratio between the standard deviation and *μtot* the mean across regions of the time-averaged rates of each region:

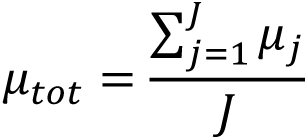

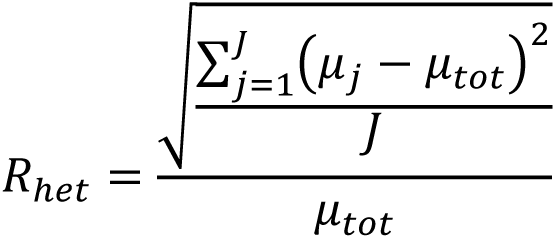

Where *μj* corresponds to the average rate of activity across time for each region *j*, *J* corresponds to the total number of regions, and *μtot* corresponds to the grand-average rate. Larger values of Rhet indicate larger heterogeneity of average rate across regions.

#### Rate variability across time

To capture the overall amplitude of oscillations in the network activity, we calculated, for each region, the CVs of rate along time, i.e., the ratio between the standard deviation and the average over time of the regional activity rate. We then took the median across regions of these CVs across regions (see Figure 2C):

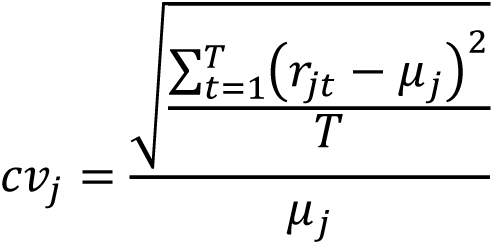

Where *rjt* corresponds to the excitatory rate of region *j* at timestep *t*, *T* corresponds to the total number of timesteps in the timeseries, and *μj* corresponds to the average rate of activity across time for each region *j*.

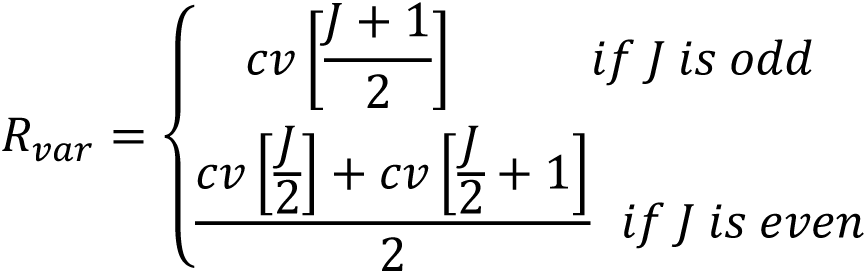

Where *cv* corresponds to the ordered list of the coefficient of variation of activity for all regions and *J* corresponds to the total number of regions in the dataset. Larger values of Rvar indicate oscillations with larger amplitude of relative variation between oscillation minima and maxima.

#### Heterogeneity of rate variability across time

Regions can be heterogeneous not only in their average rate of activity, as captured by Rhet, but may also display oscillations with different amplitudes of variation around their mean (more or less marked waves). To capture such potential heterogeneity of oscillation amplitudes we compute the CV across regions of the CVs across time of regional rates, i.e., the ratio between the standard deviation and *μcv* the average across regions (see S1):

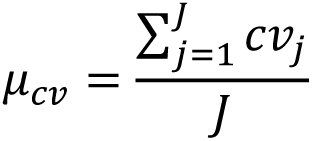

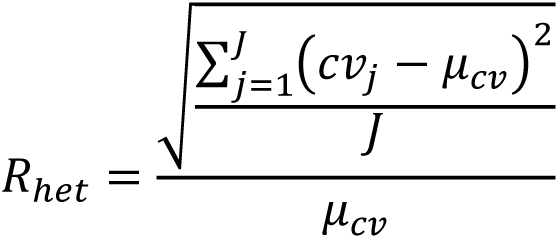

Where *cvj* corresponds to the coefficient of variation of region *j* and *J* corresponds to the total number of regions in the dataset. Larger values of Rhet indicate larger heterogeneity of oscillation amplitudes across regions.

#### Phase Locking Value (PLV)

To capture the overall level of phase synchronisation depending on the dynamic regime, we calculated the PLV among all pairs of regions (PLVall), which can disentangle phase co-fluctuation from amplitude co-fluctuation (*43*). The phase of an oscillation was obtained by performing a linear interpolation between the troughs of the oscillation to remove any effect of amplitude modulation. The troughs were detected by inverting the oscillation and using the find_peaks function from the scipy library (version 1.3.1). After extracting phases, we then evaluated the PLV for each pair of regions as:

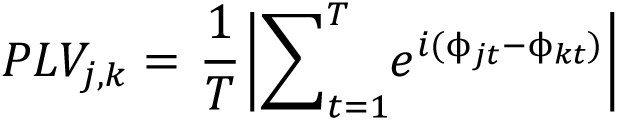

Where *ϕj/kt* corresponds to the instantaneous phase of region *j/k* at timestep *t* and *T* corresponds to the total number of timesteps in the timeseries. PLV values can range from 0 (no phase-locking) to 1 (perfect phase-locking) and is undirected as well as symmetric.

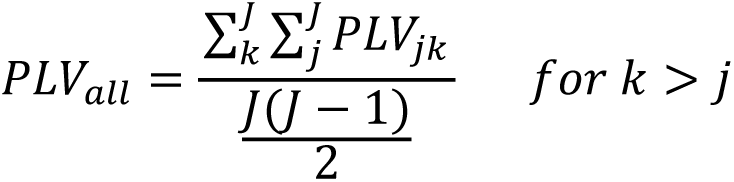

Where *PLVjk* corresponds to the PLV between region *j* and *k* and *J* corresponds to the total number of regions. Due to the symmetric nature of PLV we only calculate PLVall based on each unique PLV pair (*PLV12* = *PLV21*). An indication of the global level of synchrony over the entire network is obtained by averaging PLVs over all pairs of regions, yielding the PLVall measure. It is also possible however, that strong phase synchrony exists between a subset of regions, despite global synchrony being overall low. To capture such a potential scenario, we also evaluated the quantity PLVtop which is the average of pairwise PLVs over the top quartile of pairwise links with the highest PLV values.

### Simulating FC dynamics and extracting FC states

#### Resting-State Simulation

We used Heun’s method with an extremely precise integration timestep of 0.00005 due to the strong effect of the subsequent stimulation on the integration. The generated time-series were then down sampled, saving only one point every 200 integration steps (i.e., every 0.01 timesteps). We cut off an initial transient of 400 steps and simulated a time-series of 3000 timesteps for thirteen randomly generated initial conditions. To calculate dynamic FC measures, we used the sliding window method to obtain 140 steps windows with a 75% overlap, resulting in 1066 windows across all resting state simulation. We chose this window length in order to capture at least 50 oscillation cycles within each window, at the retained dynamic WP.

#### Time-resolved characterization of FC

In every window, we computed the value of PLV (strength of phase locking) for each pair of regions and the phase-difference at which their phase locking occurred (lag of phase locking, see later). We then compiled these PLV and phase lag values into two matrices that, together, provide a characterization of the instantaneous FC configuration.

#### Hubness and hubs

To distinguish regions that tended to participate in phase-locked coalitions i.e., being strongly phase-locked with other regions in our simulations we calculated a measure of “hubness” for each region (S2) based on time-resolved FC analyses. First, we binarized the weighted graphs described by each time resolved PLV matrices, retaining only links with PLV strength above a certain threshold. This choice naturally separated two ensembles of links naturally emerging from our simulations. Indeed, as shown by Figure S2A, the distribution of median PLV for different pairwise links has two clear peaks, one at a low PLV value of ∼0.2 and a second with a high PLV value close to ∼0.9. Both links at high and low median PLV can display a variability of the phase-lag at which phase-locking occurs (cf. Figure S2B). Retaining only links above the PLV 54th percentile threshold thus guarantees removing weakly locked links (whose phase lag is noisy and ill-defined) and only retains strongly locked links in the network which still maintain a rich variability of phase relations.

Then we calculate hubness, defined as the degree of a node (i.e., number of strong links) in each given binarized PLV window (S2C). Hubness thus provides an integer-valued metric of time-resolved centrality for a node. The most central nodes in every timeframe were denoted as “hubs”. To discriminate hub from non-hub regions in each given window we applied a second thresholding, this time not based on the PLV strength of the links, but on the hubness of the nodes. Specifically, regions were denoted as hubs if their hubness was equal to or above 21. This hubness threshold guarantees that regions that have at least one hub also have a large average PLV value (green line in Fig. SM3.2D) and, at the same time, a sufficient variability of hubs across time on average (orange line in Fig. SM3.2D). This allows for the variability of hub assignments across windows which is necessary for clustering, while for lower (higher) thresholds, variability would be non-existent, because all (no) regions are labelled as hubs across time. Note that by applying such a strong threshold, most hubness values across regions and time-windows ended up not being labelled as hubs (blue line in Fig. SM3.2D). Only roughly ∼12% of all hubness values were labelled as hubs.

#### Phase lag

To calculate the lag of phase locking between pairs of regions in each window we take the firing rate of the excitatory populations in each region and calculate their cross-correlogram, describing how the Pearson correlation between the time-series varies as a function of variable latency shift. Phase lag, normalised in the [0,1] interval (with 0 denoting in phase and 0.5 anti phase phase locking) is determined as the ratio between the latency of the first off-zero peak and the interval between the first and the second peaks (giving the average oscillation period).

Cross-correlograms were smoothed (via low-pass filtering in Fourier space) to simplify peak detection. To reduce the noise in the lag measure, we only compute phase lags between stable hub regions i.e., regions labelled as hubs at least once in every hub cluster (see *FC states: characterisation*), i.e., overall, 66 pairwise links considered for lag analyses.

#### FC states: characterisation

After characterising the FC in every time window in terms of its hubness and phase lag profiles, we grouped time-windows into discrete states using unsupervised clustering. We performed two parallel clustering, one in terms of instantaneous hubs (i.e., the 28-dimensional binary vectors of which regions are labelled as hub or not in the considered window) and a second in terms of phase lag profiles (i.e., the 66-dimensional vectors of phase-lags across all pairs of stable hub regions). We used the AgglomerativeClustering hierarchical clustering algorithm from the Scikit learn library version 0.21.3 (*98*) which starts by treating each object as a singleton cluster and then successively merges pairs of clusters based on their similarity forming a tree-based representation of their distance inter-relations. are successively merged until all clusters have been merged into one big cluster containing all objects. The resulting dendrogram is then thresholded to yield the desired number of clusters. Based on an average silhouette criterion (*99*), we chose a number of five clusters in terms of hub vectors (Fig 3A, right). Since FC states must have characteristic hub and lag profiles simultaneously, we chose four clusters in terms of phase lag vectors (Fig 3B, right). Then we constructed a contingency table between the two alternative hub and lag-based clusterings (Fig 3C, right). Hub-cluster “4” with lag-cluster “1”, hub-cluster “3” with lag-cluster “4” and hub-cluster “2” with lag cluster “2” gave rise to the three largest intersecting subsets. We thus took the time-windows belonging to these three intersection subsets and assigned to them, respectively, the FC state labels Ψ1, Ψ2, and Ψ3. All windows that did not fall inside these overlapping intersections were indistinctly termed “None”. Such construction of FC states, indeed, does not aim at attributing a state label to every possible window, but, on the contrary, at identifying sets of highly specific windows whose FC hubs and lags profiles are maximally consistent within groups and maximally distinct between groups.

To visualise similarity and differences across the FCs in different time-windows, we concatenated hubs and phase lag vectors and applied on these concatenated vectors a t-distributed stochastic neighbour embedding (t-SNE; (*46*) unsupervised algorithm for dimensionality reduction. The resulting nonlinear embedding provides a two-dimensional scatter plot in which the FC of every window is mapped to a point and in which smaller or larger distances across projected points in the 2D-plane relate to higher or lower similarities across FCs in the original (28 + 66)-dimensional space (Fig 3C, left).

#### FC states: classification

Even after applying the careful selection previously described, there is residual variability across the time-windows within each of the states Ψ1, Ψ2, and Ψ3. Furthermore, when applying stimulation, the system generates previously unseen FCs for which there is no state assignment available from unsupervised clustering. It is thus necessary to identify FC windows maximally representative of the specific state to which they belong (state “prototypes”) and to construct a supervised classifier which is able to assign a state label to unseen windows based on their degrees of similarity to the different state prototypes. To quantify how representative of an FC state each time window is, we summed the two silhouette scores (*99*) of the window obtained independently for hub-based and lag-based clusters. The silhouette score values range between 1 (best) and -1 (worst) and quantify how similar a sample is to its own cluster relative to how similar it is to other clusters. For each state, we thus select 100 windows with the highest *typicality* (additive silhouette score) per state.

To identify the FC states in novel post-stimulation samples, we constructed a *k-*nearest neighbours classifier using these 100 prototypes per state as training set. This classifier identifies the *k* nearest training set samples for a novel sample and assigns the novel sample the state label to which the majority of its nearest neighbours belong to. A choice of k = 50 guaranteed the best generalisation performance over the remaining non-prototypical resting state windows for which a ground truth state label was available.

### Stimulation

#### Stimulation Simulation

We applied a punctual single pulse stimulation to emulate Dirac’s delta applied to both the inhibitory and excitatory populations of the stimulated region and simulated 500 timesteps post stimulation. Stimulation was systematically applied at different phases of the ongoing oscillation cycle, from 0 to 2π in 10 intermediate steps. We used a stimulation intensity of 300 (2 magnitudes larger than the adopted background drive Iext) and performed stimulation experiments in six different regions (V1, DP, TEO, 2, 8B, STPr), and three different FC states (Ψ1, Ψ2, Ψ3). For each FC state, we ran simulated stimulation experiments in five different time-windows (the five prototype window with the topmost values of combined silhouette, see previous section) and in five subsequent oscillation cycles, resulting in 25 simulations samples in total for each combination of region, phase and state, provided that they lay 500 simulation steps before the end of the resting state simulation. Due to computational tractability we had to make a subselection of regions to stimulate in our simulation. The rationale for choosing the six regions we considered was the following: We covered most cortical areas as TEO is in the visual area, STPr is in the motor area, DP is in the parietal area, and 8B is in the prefrontal area; furthermore, investigate regions with two extreme roles with respect to FC topology we also stimulated V1, which has the lowest probability serving as hub and 2, which has the highest probability.

#### Phase Shifting Portraits

We obtained the phase shifting portrait of each stimulation simulation by calculating the phase difference for each stimulated phase between stimulated and unstimulated time-series at each of the 500 timesteps post-stimulation and wrapped the difference in a range between - 0.5π and + 0.5π (as phase shifts of -0.7π and + 0.3π functionally correspond; S5). To compute the non-stimulated phase time-series for every stimulation simulation, we ran a parallel simulation which was identical in all aspects except the stimulation intensity which was set to zero.

Group-level phase shifting portraits (Figure 4) were calculated for each parameter combination of stimulated region, phase and FC state. We obtained the group-level phase shifting portraits by averaging the phase shifting portraits of individual stimulation simulations over the 25 samples of stimulation with the same region, phase and FC state. Averages had to be performed on the complex plane as the quantities to be averaged are phase differences on the unit cycle.

#### State Morphing: Phase Response Curve (PRC)

To obtain the PRC of a given region (Figure 5C), we started by averaging the phase difference between stimulated and unstimulated simulations, in each individual stimulation simulation, over the last 140 steps window to discard initial transients induced by the stimulation pulse. Then, prior to averaging these phase shifts across samples we excluded samples where the classified FC state of the post-stimulation window was not the same as the FC state at the time of stimulation. In this way, we could guarantee that the retained stimulation experiments were inducing only state morphing, and thus evaluate state-specific PRCs. Averages of phase shifts are performed once again on the complex plane. Since stimulating a region induces highly nonlocal phase shifting even in distant regions, we had to evaluate an individual PRC for each affected region in each parameter combination of phase, state and region of stimulation.

#### State Switching

To capture the probability difference of state switching due to stimulation we first applied the FC classifier (see *FC states: classification*) to FC from the same post-stimulation window used for PRC estimation (i.e., 140 timesteps post-stimulation), and using the 25 samples to estimating the post-stimulation probability of state switching for each parameter combination. As state switching can also occur spontaneously, we also computed this probability for the matching unstimulated simulation. We then subtracted the unstimulated from the stimulated probabilities of switching to obtain the probability difference of stimulation-induced state switching beyond possible spontaneous state switching in the unstimulated time-series for each combination of phase-, region and state-dependent measures (Figure 5A, D). For positive values of the probability difference the stimulation is inducing state switching, whereas negative values reflect a suppression of possibly natural state switching. Note that we perform stimulation in highly prototypical (and thus stable) windows, so that the unstimulated probabilities of switching were very small for Ψ1 and Ψ3 (and consistently at 0.2 for Ψ2).

#### Prediction

To predict stimulation-dependent phase-shifting in the last window of the post stimulation timeseries we use a Random Forest Regression (RFR) algorithm, implemented within the scikit-learn library (version 0.21.3; (*98*)). In such an approach, various decision trees are constructed in which the final leaves correspond to (discretized) predictions of the obtained phase-shift and branches to follow are selected based on the values of the chosen input features. The final prediction is obtained averaging over a “forest” of many randomised trees. We used a maximal depth of 5 levels of branching for trees in the forest. We trained an independent RFR for each pair formed by a stimulated region (one of 6 possible: V1, DP, TEO, 2, 8B, STPr) and a region where effects were observed (one of 28 possible), i.e., 168 classifiers to train (for each choice of the set of input features). Since the target output is a phase-shift over the unit polar circle, but RFRs operate on the real axis, we transformed ΔΦ into a complex number exp^iΔΦ^ and used the real and imaginary part as the two dimensions of the output vector. Predictors included FC state-ignorant measures (phase of stimulation of the stimulated region; outgoing structural connectivity of the stimulated region SCloc) and FC state-aware measures (FC state label, PLV, PLVlocal, Lag, Laglocal). The phase of stimulation was a 2 × 750 vector containing the real and imaginary phase of stimulation in radians at the stimulated region; the structural connectivity was a 29 × 750 vector consisting of the vector of outgoing structural connectivity of the stimulated region; the FC state was a 1 × 750 vector with the identity of the stimulated FC state identity; the PLV was a 406 × 750 vector containing the upper triangle of the PLV matrix in the corresponding stimulated window; PLVlocal was a 8 (V1), 13 (TEO), 24 (STPr), 17 (DP), 16 (2), 18 (8B) × 750 vector containing the PLV between the stimulated region and the regions it structurally projected to; the Lag was a 406 × 750 vector containing the upper triangle of the lag matrix in the corresponding stimulated window; Laglocal was a 8 (V1), 13 (TEO), 24 (STPr), 17 (DP), 16 (2), 18 (8B) × 750 vector containing the PLV between the stimulated region and the regions it structurally projected to. To explore how various parameter combinations affect the prediction accuracy we concatenated the chosen predictors along the non-sample dimension so that we ended up with a *n* × 750 input vector. To make sure our results could generalise to novel data we use ten-fold cross-validation with a training size of 70 %. Furthermore, we also stratified according to FC state, as classifiers may not be able to recognize classes that they have not been trained on.

## Acknowledgements

We thank Giovanni Rabuffo, and Jan Fousek for the technical support in creating the model within the TVB software.

## Author contributions

Conceptualization: BG, DB

Methodology: SBS, DB

Investigation: SBS, DB

Visualisation: SBS

Supervision: MG, DB

Writing—original draft: SBS

Writing—review & editing: MG, DB

## Data and materials availability

All data needed to evaluate the conclusions in the paper are present in the paper and/or the Supplementary Materials.

## Supplementary Figures

**Figure S1.**
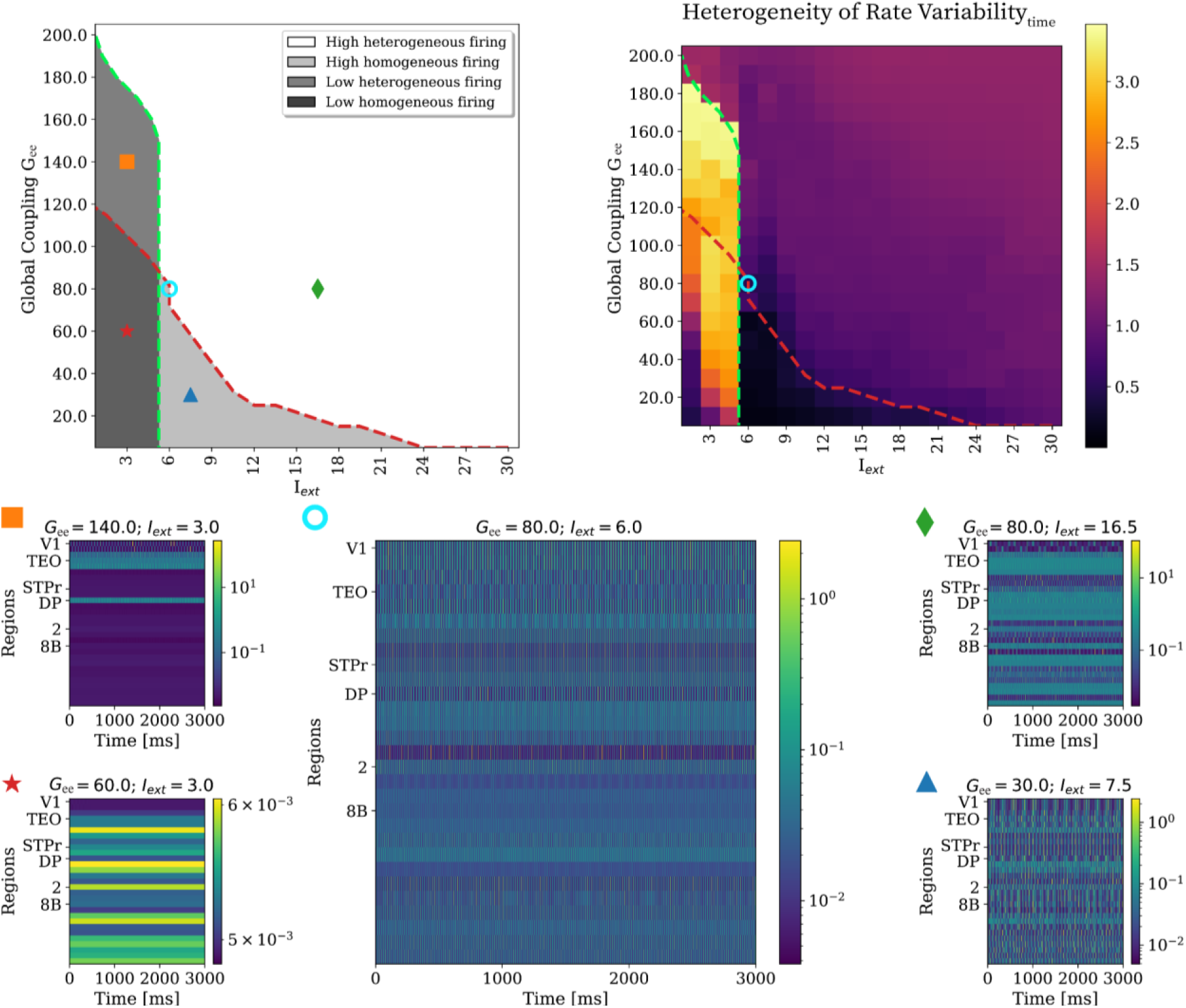
Example time series of regimes. Top left: Dynamical regimes for values of global excitatory-to-excitatory coupling Gee between 10 and 200 versus values of the local drive to the excitatory population Iext between 1.5 and 30. The red dashed line divides regimes whose activity is homogeneous across regions (below) from regimes whose activity is heterogeneous across regions (above). The green dashed line divides regimes who are highly active (right) from regimes who are not as active (left). Symbols mark examples of the four categorical dynamical regimes: blue triangle (homogeneously low firing rate; dark grey), orange square (homogeneously high firing rate; light grey,), green rhombus (heterogeneously low firing rate; intermediate grey) red star (heterogeneously high firing rate; white). The black dashed circle represents the chosen WP (Gee= 80.0, Iext = 6.0). Top right: Rate heterogeneity across times for the same parameter ranges of Gee and Iext as in the top left. To capture how heterogeneous the variation of amplitudes is across regions, we calculate the coefficient of variation across time for each region and calculate the coefficient of variation of this value across regions (see *Methods* for more details). Bottom: Example time series simulated for 300000 timesteps for each parameter combination corresponding to each symbol shown in the top left. Annotated regions correspond to the stimulated regions.

**Figure S2.**
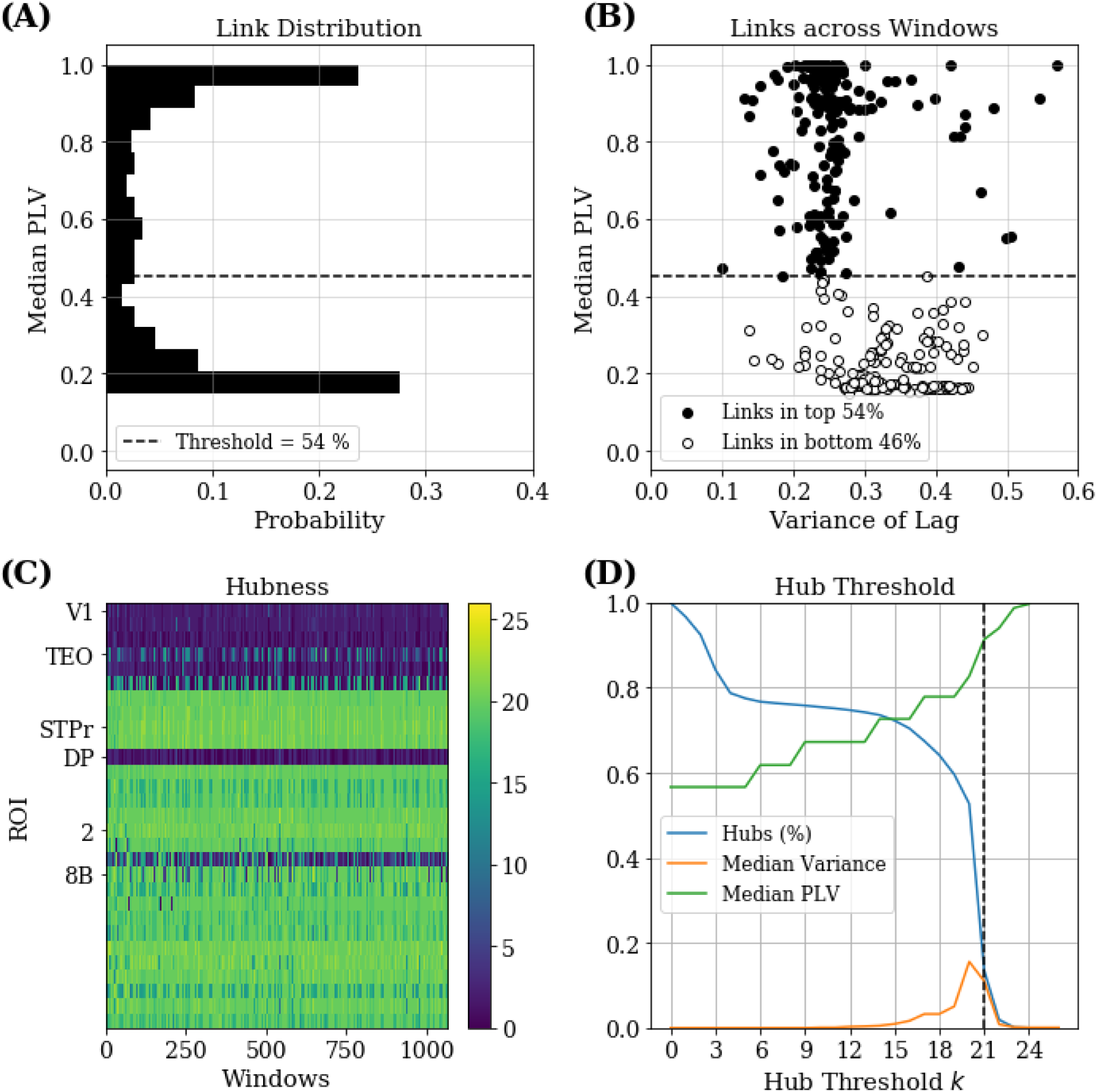
Pipeline for calculating hub identity. 1066 windows were obtained with the sliding window approach (window length = 140 timesteps, overlap = 75%) from resting-state time-series with 300000 timesteps for 13 different initial conditions with the parameters corresponding to the chosen WP (Gee= 80.0, Iext = 6.0). (A) Probability distribution of average PLV for each link between two regions across all 1066 windows. Dashed line marking 54% threshold for dividing links into links with high phase-locking (1) or links with low phase-locking (0). (B) Average PLV of each link calculated across windows plotted against the variability of lag of each link across windows. The dashed line marks the threshold of 54th percentile of PLV values separating the white point cloud with a large variability in lag suggestive of noise from the dark point cloud with a smaller variability in lag. (C) Time-resolved hubness. Hubness corresponds to the sum of high PLV links for each of the 28 regions in the 1066 windows, based on previous binarization with a threshold of 54%. Annotated regions correspond to the later stimulated regions. (D) Effect of chosen threshold for hub detection in hubness on percentage of hubs overall (blue line), average PLV of regions with hubs (green line), and variability of hubs over windows (orange line). Chosen threshold for hub identity in dashed black line (*k* = 21).

**Figure S3.**
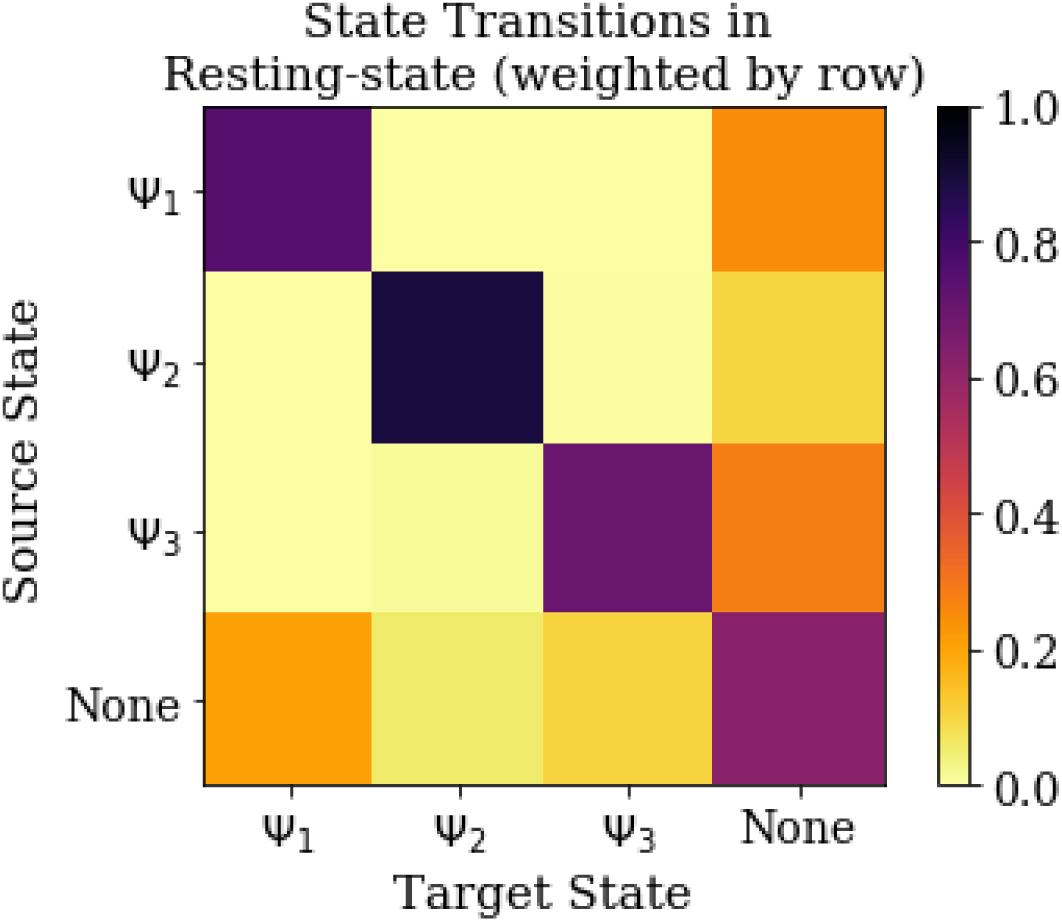
Temporal stability of state trajectories. FC states correspond to three largest intersecting labels extracted from the independent clustering of hub identity and phase lag. Ψ1 belongs to hub-cluster “4” with lag-cluster “1”, Ψ2 belongs to hub-cluster “3” with lag-cluster “4” and Ψ3 belongs to hub-cluster “2” with lag cluster “2” gave rise to the three largest intersecting subsets. All windows corresponding to the “None” state do not fall under the joined hub identity and lag definition of states Ψ1, Ψ2, Ψ3. See grey dots in the t-SNE projection and grey squares in the confusion matrix in Fig 3C. The source state represents which of the four groups (Ψ1, Ψ2, Ψ3, None) a window belongs to at time point t. The target state represents which of the four groups a window belongs to at time point t + 1. Since the numbers for each of the four groups are not balanced, we normalise the matrix by row. The probabilities on the diagonal are high, meaning that it is very likely to be in the same group at time point t + 1 as at time point 1 suggesting a high temporal stability of the states.

**Figure S4.**
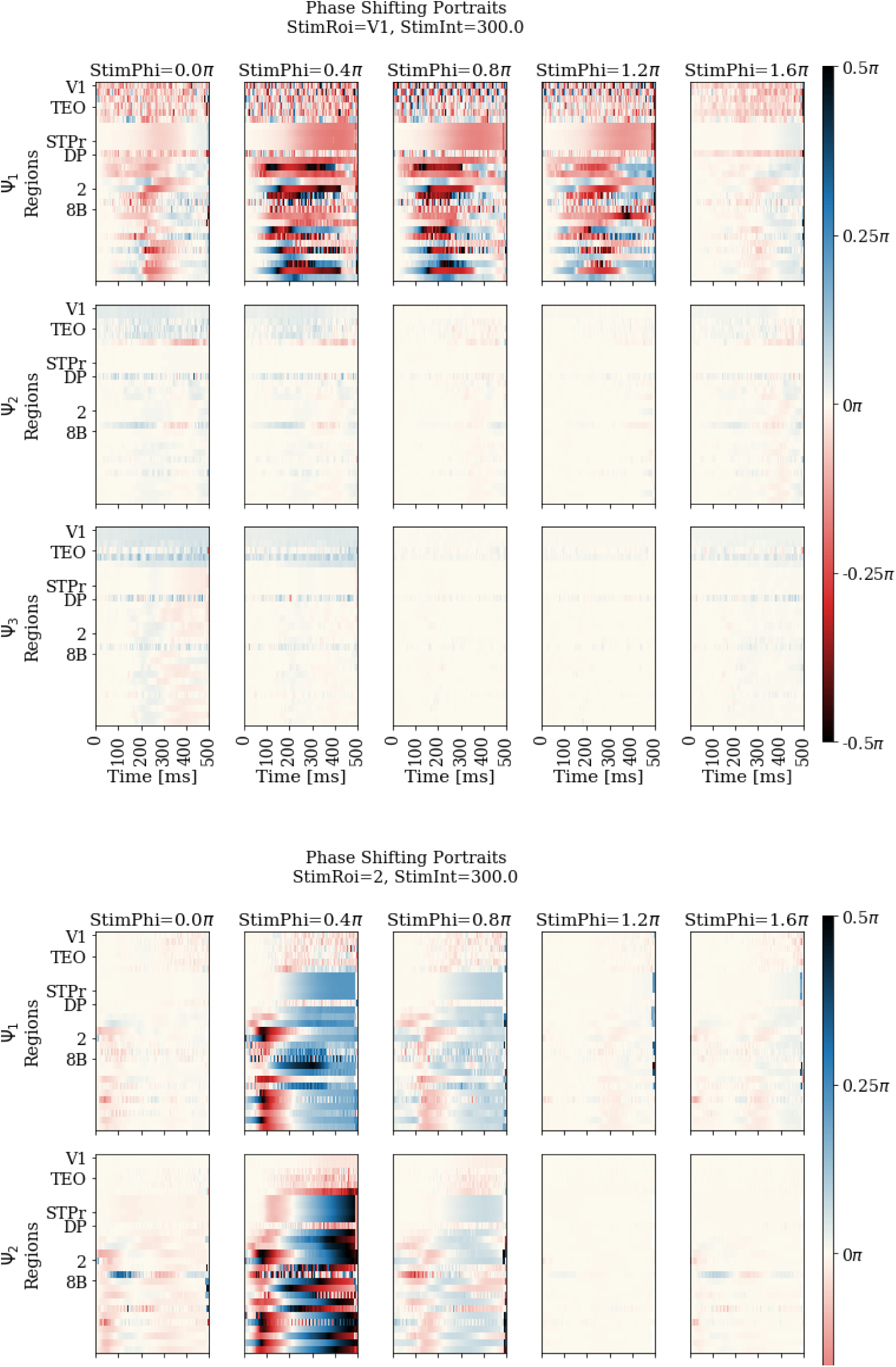
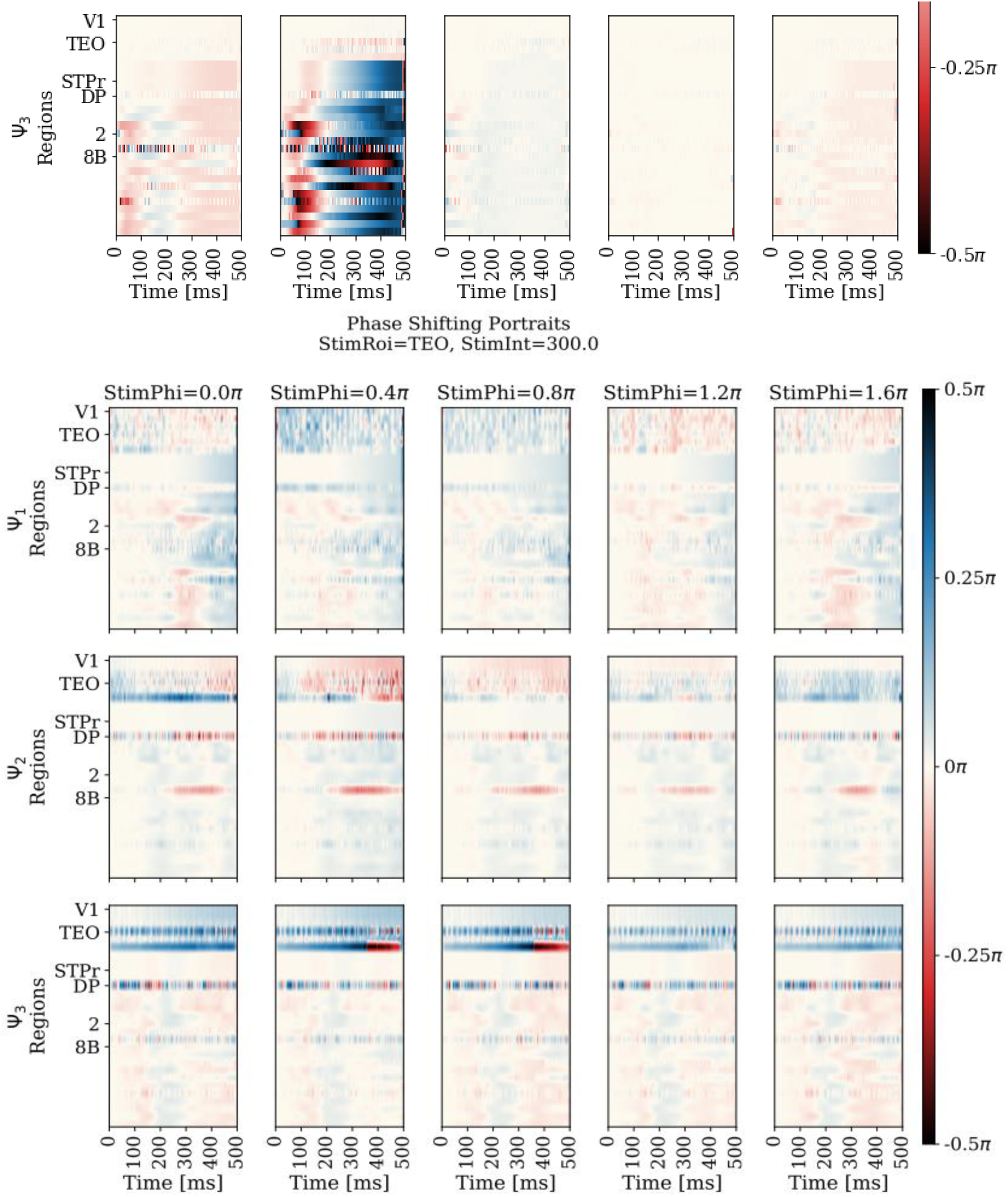
Phase shifting portraits (Δϕ) for V1, TEO, and 2. We perform identical stimulation experiments in the five most typical windows of each FC state and in five subsequent oscillatory cycles in each typical window equalling 25 stimulation experiments for each parameter combination of stimulated phase, FC state, and region. First, we calculate the phase difference between stimulated and unstimulated time series for 50000 timesteps post stimulation and wrapped the difference in a range between - 0.5π and + 0.5π. Then, for each portrait the angle of the complex average is calculated across the 25 samples for each parameter combination of stimulated phase, FC state, and region. The phase shifting portraits are again wrapped in a range between - 0.5π and + 0.5π. Labelled regions are the regions stimulated in the stimulation simulation. Effects of phase, FC state, and region on phase shifting can be multifactorial and conditionally dependent.

**Figure S5.**
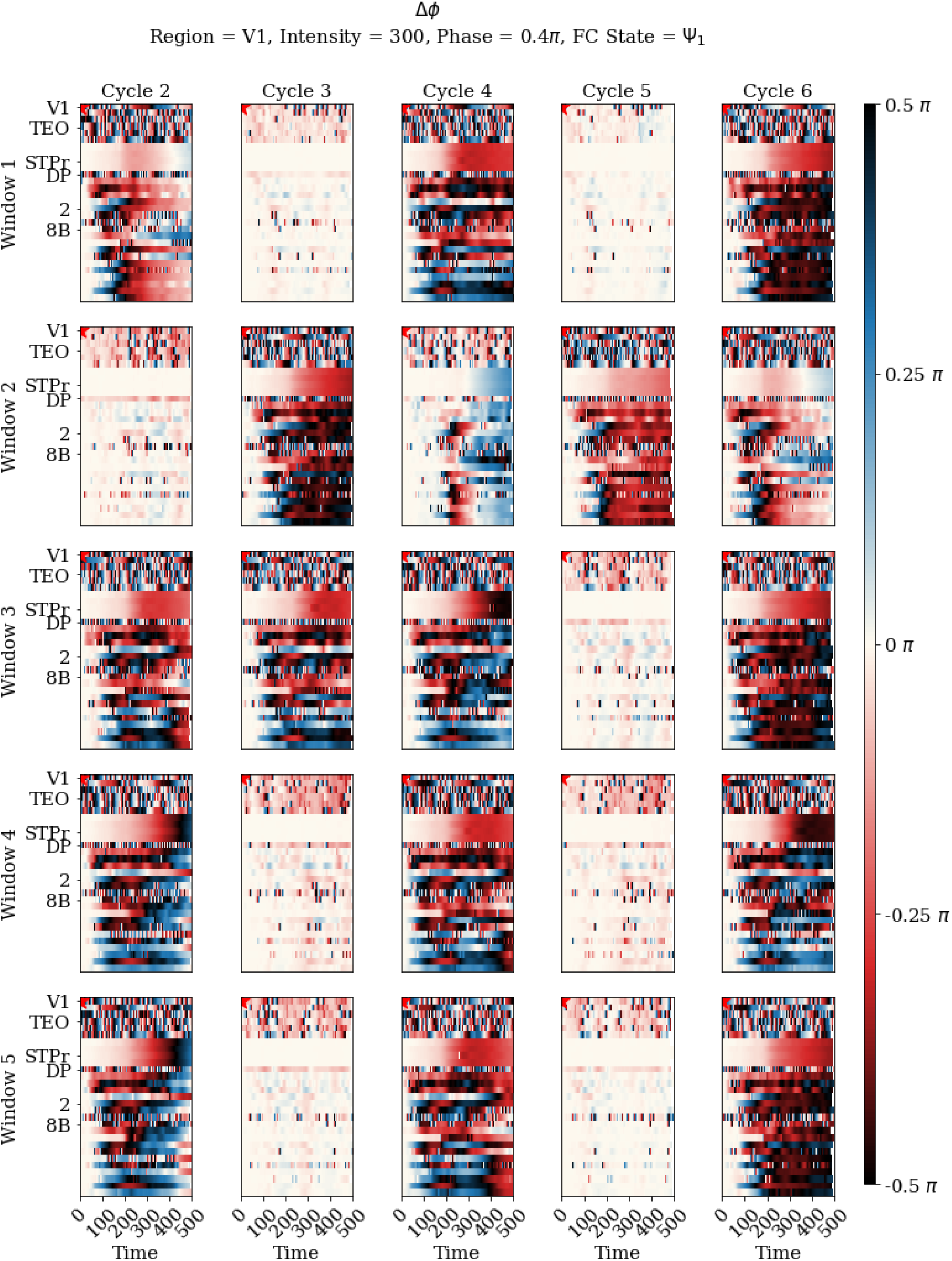
Example of variability across 25 samples for stimulation phase 0.2π, FC state Ψ1 and region V1. The Δϕ is shown for each of the 25 samples for stimulation phase 0.4π, FC state Ψ1 and region V1. Windows correspond to the five most typical windows for FC state Ψ1 and cycles refer to the oscillatory cycle of each window in which stimulation was applied. Δϕ is calculated by obtaining the phase difference between stimulated and unstimulated time series for 50000 timesteps post stimulation and wrapping the difference in a range between - 0.5π and + 0.5π. Labelled regions correspond to all regions that are stimulated in general, not in this example, specifically. Samples for some parameter combinations can show a tremendous amount of variability.

**Figure S6.**
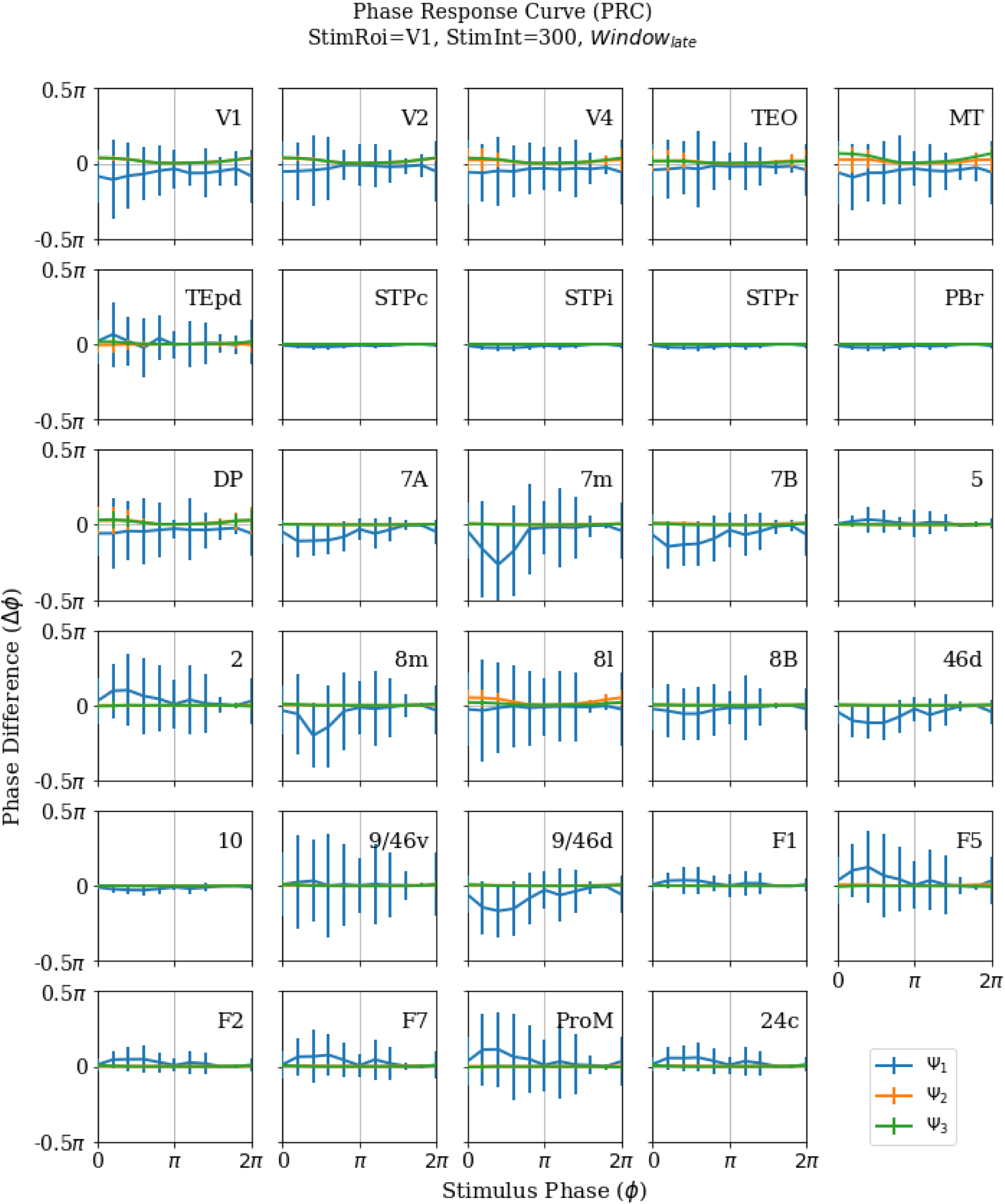
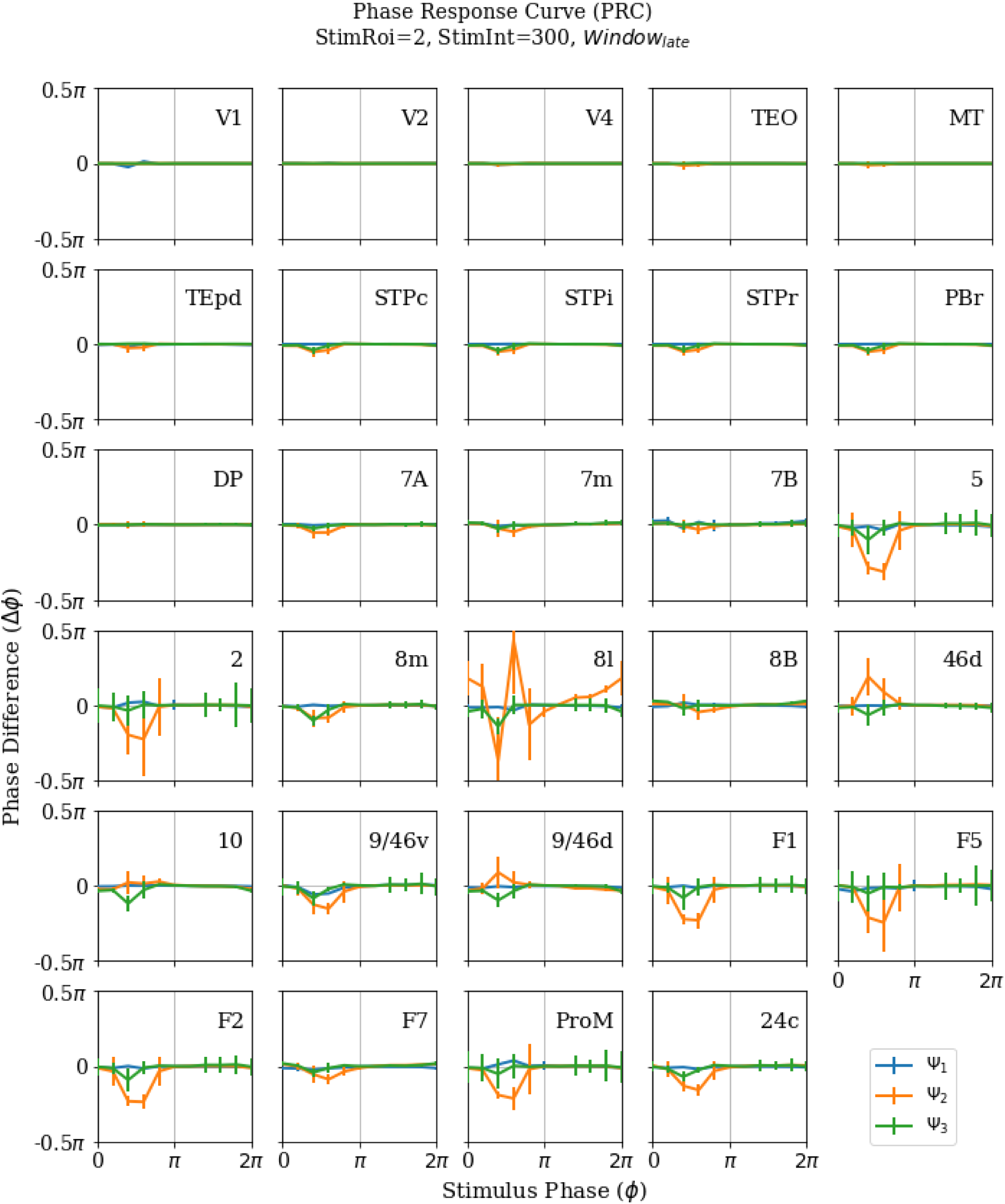
PRC in state morphing for regions V1 and 2. PRCs are calculated for every affected region independently for each parameter combination of FC state and stimulated region. Of the 25 original trials only, those are selected that remain in the same FC state at stimulation as at the most posterior window i.e., those trials that only undergo state morphing. In those trials, we calculate the phase difference between unstimulated and stimulated trials in the most posterior 1400 timestep window of the 50000 post-stimulation time series. Then, we calculate the angle of the complex average for each parameter combination of stimulated phase, FC state, and stimulated region based on the state morphing subset. Vertical lines in FC state colours show the standard deviation. The PRCs for regions STPr, 8B, and DP remain nearly entirely flat irrespective of region and FC state, whereas the PRCs in Ψ1 (Ψ3) show a strong variability when stimulating region V1 (TEO).

**Figure S7.**
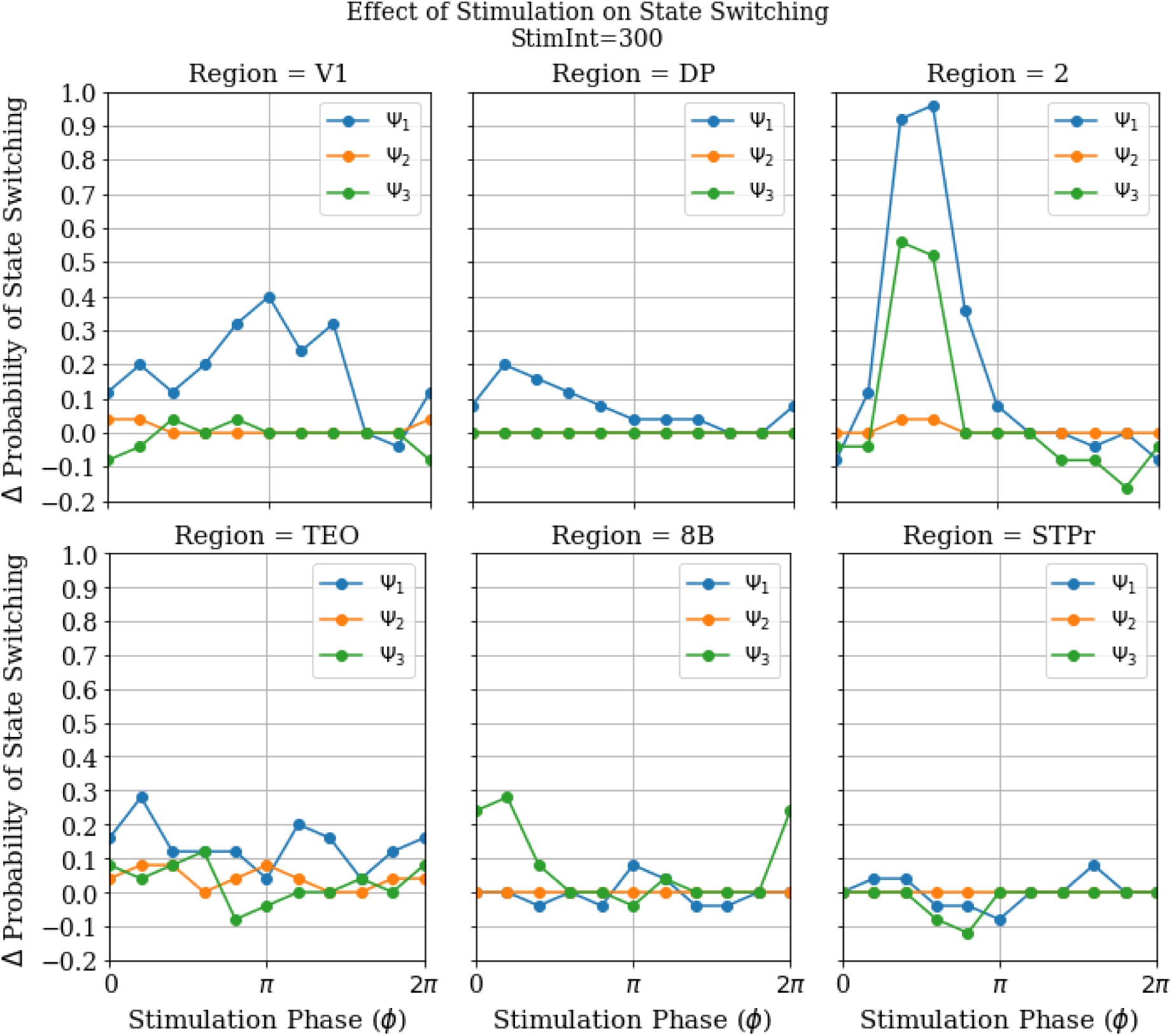
Probability difference of state switching for V1, TEO, STPr, DP, 2, and 8BP. The probability difference of state switching is calculated for each parameter combination of stimulated phase, FC state, and stimulated region. First, the FC state in the most posterior 1400 window in the 500000 timesteps post-stimulation is quantified with machine learning for all 25 samples of each parameter combination. Then, the probability of switching to a different FC state is calculated for the stimulated trials and subtracted by the probability of switching to a different FC state for the unstimulated trials. The state switching probability differences of regions TEO, STPr, DP, and 8B remain below 0.3 irrespective of FC state or phase of stimulation. There is a strong phase-dependent state switching probability for FC state Ψ1 and Ψ3 when stimulating region 2 at 0.2π and 0.3π.

**Figure S8.**
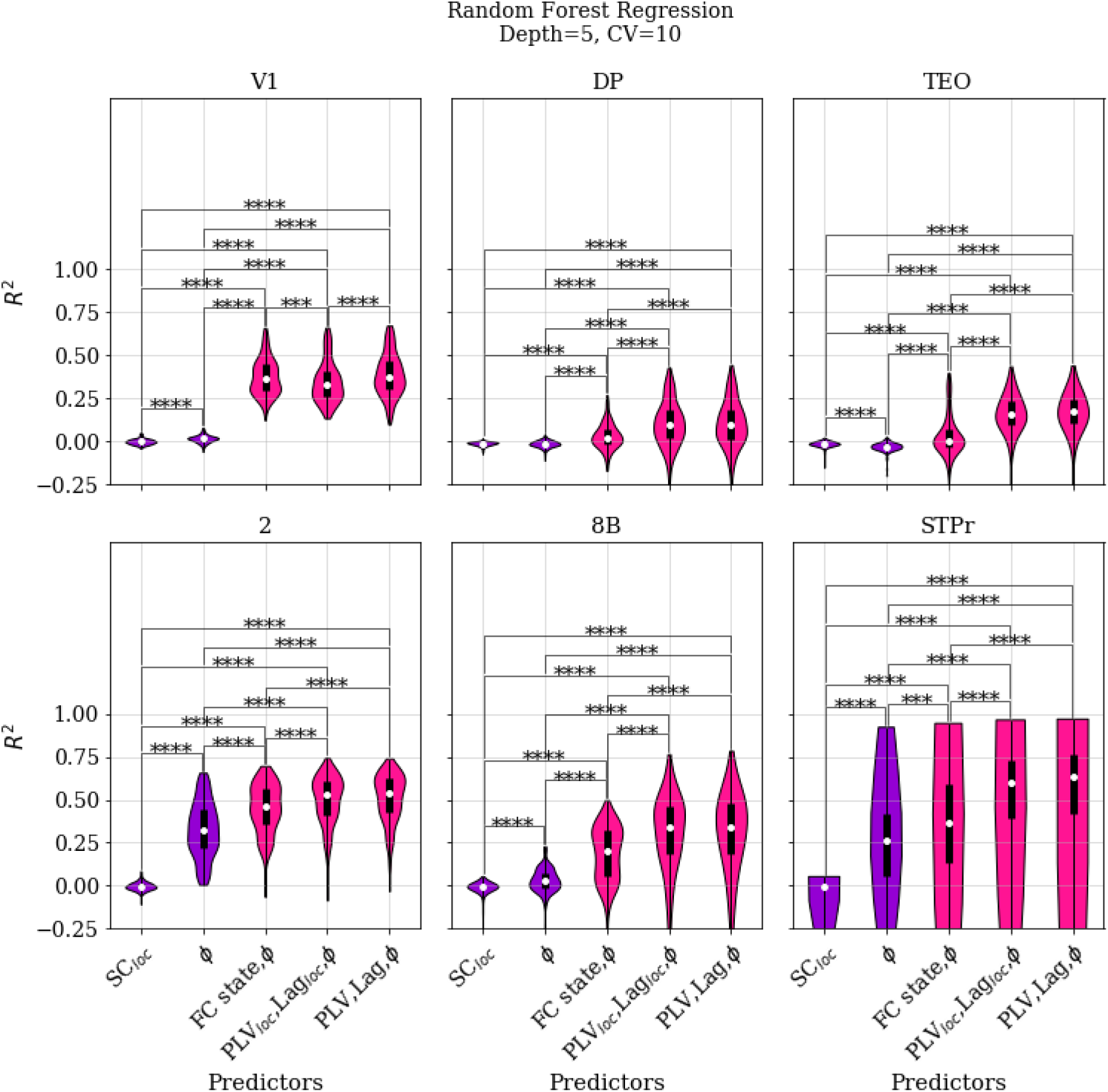
Performance of random forest regression in predicting stimulation-induced phase-shifting for V1, TEO, STPr, DP, 2, and 8B. The classification aims to predict the stimulation-dependent phase difference between stimulated and unstimulated time series in the most posterior 1400 window in the 500000 timesteps post-stimulation with a Random Forest Regression (RFR) algorithm. 168 independent classifiers are trained for each combination of the 6 stimulated regions and 28 affected regions. Each combination contained 750 trials in total across the 10 stimulation phases, the 3 FC states, and the 25 samples for each FC state. We used 10-fold cross validation (training size = 70 %) to show how generalizable the information is which was learned by the classifier leading. The performance of the classifier improved significantly for all stimulated regions when state-aware predictors (PLV, PLVlocal, Lag or Laglocal) were used. Interestingly, using FC state and stimulated phase did not significantly outperform SCloc and stimulated phase for DP, TEO, 2, 8B, and STPr. Each violin plot reflects the test performance for 10 CV folds for all of the 29 affected regions (290 points in total). In each violin plot the white dot marks the median and the thick black line marks the inner quartiles. Significance was tested with the Mann-Whitney U test and significance levels are indicated as follows: not drawn for p ≥ 0.05; * for p < 0.05; ** for p < 0.01; *** for p < 0.001; **** for p < 0.0001. Bonferroni correction for multiple comparisons was applied.

**Figure S9.**
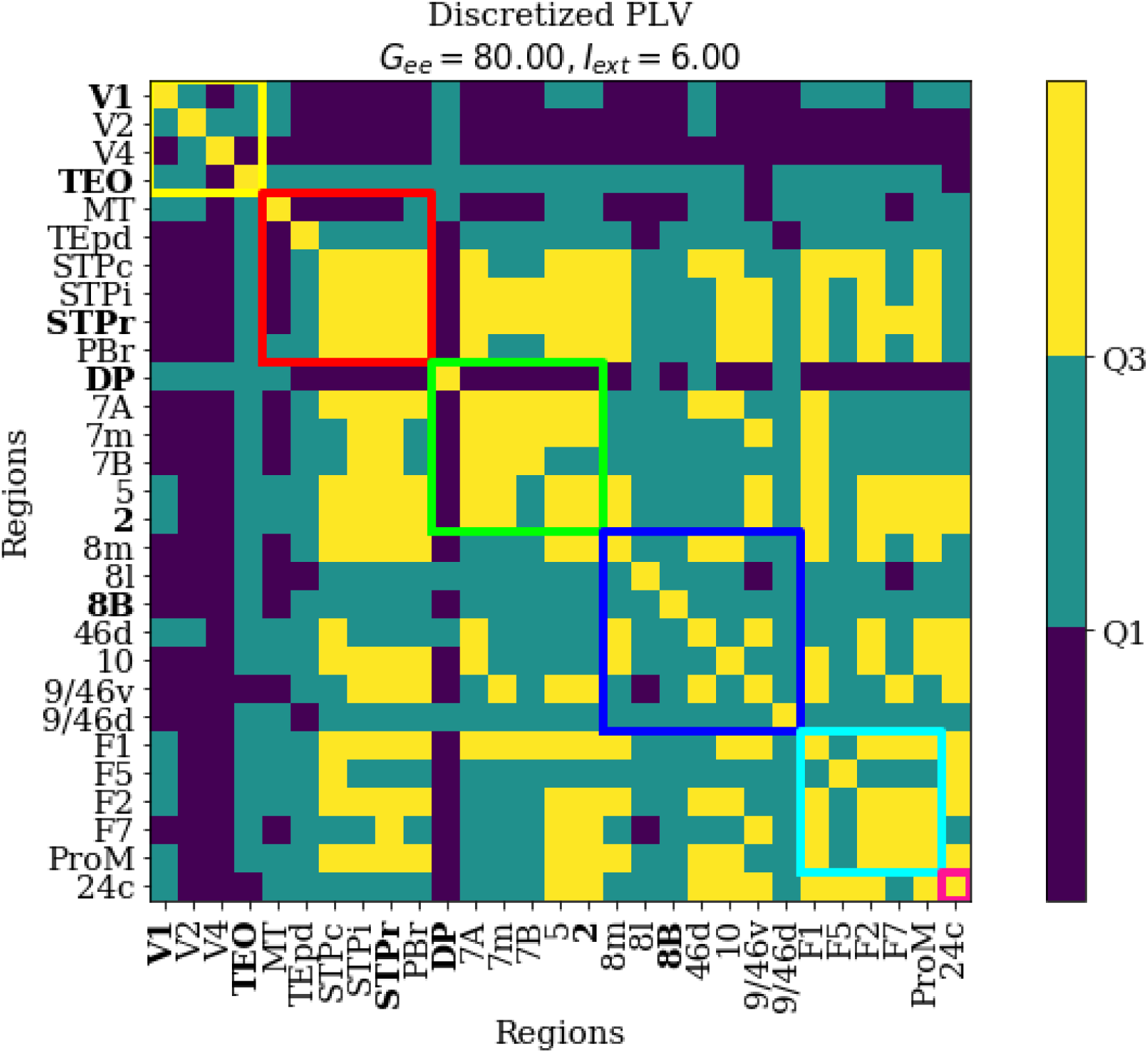
Discretized PLVall for chosen WP. Phase-Locking Values were calculated across the entire resting-state time-series for a single random initial condition for Gee = 80 and Iext = 6 i.e., the chosen WP. Dark blue represents the values that lie inside of the bottom quartile, teal represents the centre 50 percentile of values and yellow represents the top quartile of PLV values. Interareal connectivity is marked by squares (yellow: visual area, red: motor area, green: parietal area, dark blue: prefrontal area, light blue: frontal area, pink: limbic area). Labelled regions correspond to all the regions that were stimulated in the stimulation simulation.

**Figure S10.**
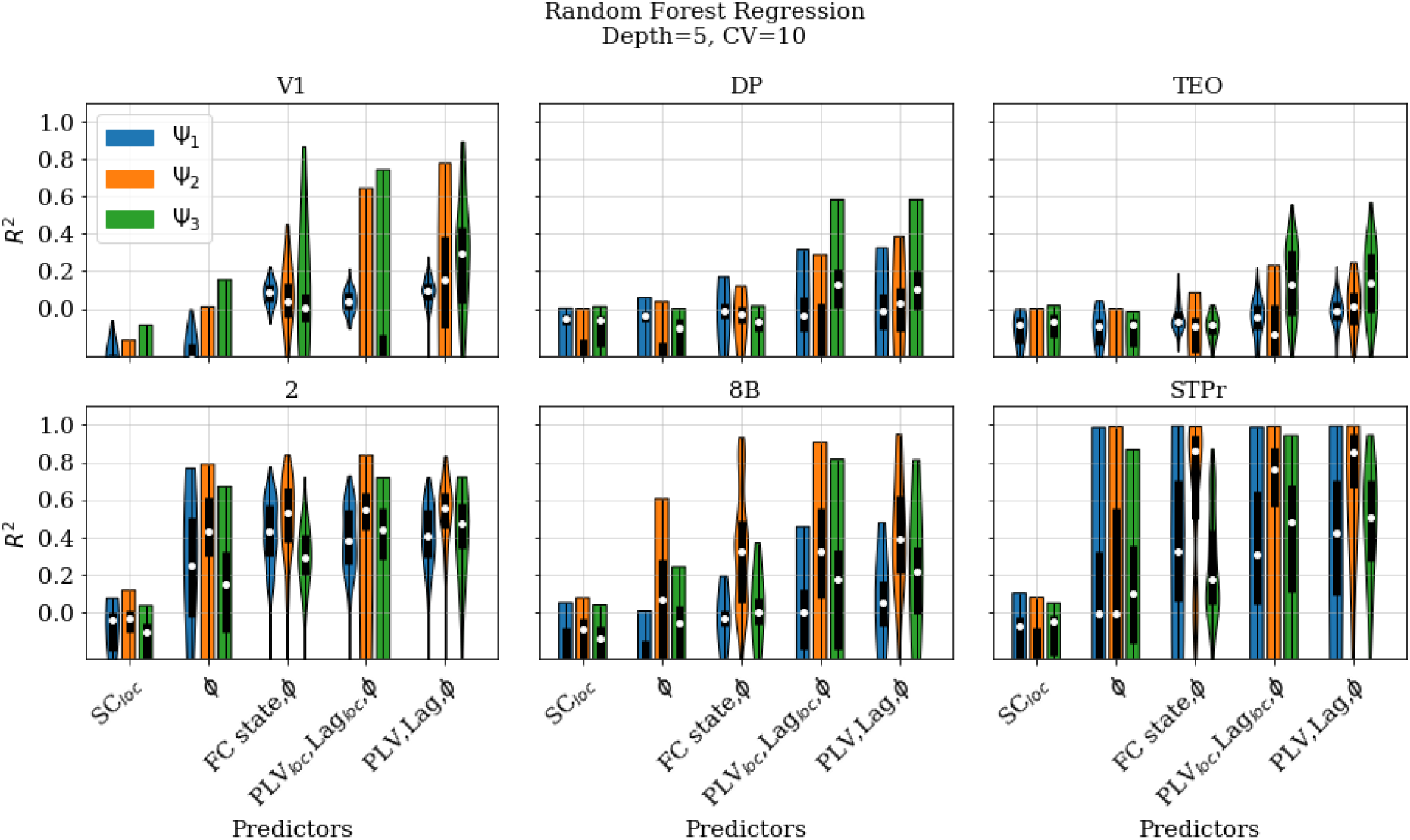
Performance of random forest regression in predicting stimulation-induced phase-shifting for V1, TEO, STPr, DP, 2, and 8B split according to FC state. The classification aims to predict the stimulation-dependent phase difference between stimulated and unstimulated time series in the most posterior 1400 window in the 500000 timesteps post-stimulation with a Random Forest Regression (RFR) algorithm. 168 independent classifiers are trained for each combination of the 6 stimulated regions and 28 affected regions. Each combination contained 750 trials in total across the 10 stimulation phases, the 3 FC states, and the 25 samples for each FC state. We used 10-fold cross validation (training size = 70 %) to show how generalizable the information is which was learned by the classifier. The performance of the classifier improved significantly for all stimulated regions when state-aware predictors (FC state, PLV, PLVlocal, Lag or Laglocal) were used. In each violin plot the white dot marks the median and the thick black line marks the inner quartiles. Performance was split across FC states. The reduction in performance could not be attributed to the systematic underperformance of a single FC state but was spread over all FC states.

